# Diverse signatures of convergent evolution in cacti-associated yeasts

**DOI:** 10.1101/2023.09.14.557833

**Authors:** Carla Gonçalves, Marie-Claire Harrison, Jacob L. Steenwyk, Dana A. Opulente, Abigail L. LaBella, John F. Wolters, Xiaofan Zhou, Xing-Xing Shen, Marizeth Groenewald, Chris Todd Hittinger, Antonis Rokas

## Abstract

Many distantly related organisms have convergently evolved traits and lifestyles that enable them to live in similar ecological environments. However, the extent of phenotypic convergence evolving through the same or distinct genetic trajectories remains an open question. Here, we leverage a comprehensive dataset of genomic and phenotypic data from 1,049 yeast species in the subphylum Saccharomycotina (Kingdom Fungi, Phylum Ascomycota) to explore signatures of convergent evolution in cactophilic yeasts, ecological specialists associated with cacti. We inferred that the ecological association of yeasts with cacti arose independently ∼17 times. Using machine-learning, we further found that cactophily can be predicted with 76% accuracy from functional genomic and phenotypic data. The most informative feature for predicting cactophily was thermotolerance, which is likely associated with duplication and altered evolutionary rates of genes impacting the cell envelope in several cactophilic lineages. We also identified horizontal gene transfer and duplication events of plant cell wall-degrading enzymes in distantly related cactophilic clades, suggesting that putatively adaptive traits evolved through disparate molecular mechanisms. Remarkably, multiple cactophilic lineages and their close relatives are emerging human opportunistic pathogens, suggesting that the cactophilic lifestyle—and perhaps more generally lifestyles favoring thermotolerance—may preadapt yeasts to cause human disease. This work underscores the potential of a multifaceted approach involving high throughput genomic and phenotypic data to shed light onto ecological adaptation and highlights how convergent evolution to wild environments could facilitate the transition to human pathogenicity.

## Introduction

Convergent evolution, the repeated evolution of similar traits among distantly related taxa, is ubiquitous in nature and has been documented across all domains of life ^1–3^. Convergence typically arises when organisms occupy similar ecological niches or encounter similar conditions and selective pressures; facing similar selective pressures, organisms from distinct lineages often evolve similar adaptations.

Independently evolved phenotypes often share the same genetic underpinnings (parallel molecular evolution) ^4–7^ such as similar mutations in specific genes ^4,5,8^, but can also arise through distinct molecular paths and by distinct evolutionary mechanisms ^9,10^, such as copy number variation ^11–13^, gene losses ^14–16^, or gene gains (e.g., horizontal gene transfer or HGT) ^17,18^. Molecular signatures of convergence can also be inferred from independent shifts in overall evolutionary rates ^19^ and examined at higher hierarchical levels of molecular organization, such as functions or pathways ^20^. For instance, comparing rates of evolution across distantly related animal lineages could pinpoint convergent slowly evolving genes involved in adaptive functions^21^ or convergent rapidly evolving genes indicating parallel relaxed constraints acting on dispensable functions ^22^. Parallel molecular changes are common across all domains of life ^9^, but their occurrence can be reduced by mutational epistasis or the polygenic nature of some phenotypic traits ^10,23,24^, particularly when studying convergence in distantly related organisms.

Fungi exhibit very high levels of evolutionary sequence divergence ^25^; the amino acid sequence divergence between the baker’s yeast *Saccharomyces cerevisiae* and the human commensal and opportunistic pathogen *Candida albicans*, both members of subphylum Saccharomycotina (one of the three subphyla in Ascomycota, which is one of the more than one dozen fungal phyla) is comparable to the divergence between humans and sponges ^26^. Due to their very diverse genetic makeups, convergent phenotypes arising in fungi might involve distinct genetic determinants and/or mechanisms, including HGT ^27,28^, a far less common mechanism among animals (but see ^29,30^).

Saccharomycotina yeasts are ecologically diverse, occupy diverse ecosystems ^31^, and vary considerably in their degree of ecological specialisation ranging from cosmopolitan generalists to ecological specialists. For instance, *Sugiyamaella* yeasts are mostly isolated from insects ^32^ and most *Tortispora* species have been almost exclusively found in association with cacti plants ^33^. The cactus environment accommodates numerous yeast species rarely found in other niches ^34–37^. Moreover, cactophilic yeasts are part of a model ecological system involving the tripartite relationship between cactus, yeast, and *Drosophila* ^34,36,38,39^. Cactophilic yeasts use necrotic tissues of cacti as substrates for growth ^40^ while serving as a food source to cactophilic *Drosophila*. Cactophilic flies (and other insects) play, in turn, a crucial role in the yeast’s life cycle by acting as vectors ^34^.

In *Drosophila*, the adoption of cacti as breeding and feeding sites evolved ∼16-21 million years ago (Mya) and is considered one of the most extensive and successful ecological transitions within the genus ^41^. Cactophilic *Drosophila* thrive across a wide range of cacti species that differ in the profiles of toxic metabolites they produce – *Opuntia* species, commonly called prickly pear cactus, generally contain fewer toxic metabolites than columnar cacti species ^40^ and are likely the ancestral hosts ^41^. The distinctive characteristics of the cacti environment ^42^ seemingly selected for adaptive traits across cactophilic *Drosophila*, such as high resistance to heat, desiccation, and toxic alkaloid compounds produced by certain types of cacti ^43–48^. These traits are likely associated with several genomic signatures (e.g., positive selection, gene duplications, gene gains) impacting multiple functions, such as water preservation or detoxification ^45–50^.

Contrasting with *Drosophila*, where cactophily is largely found within the monophyletic *D. repleta* group ^41^, molecular phylogenetic analyses revealed that cactophilic yeasts belong to phylogenetically distinct clades, indicating that association with cacti evolved multiple times independently in the Saccharomycotina ^37^. While relevant ecological and physiological information of cacti-associated yeasts is available ^35–37,39^, the genetic changes that facilitated the convergent evolution of multiple yeast lineages to the cacti environment are unknown.

Benefiting from the wealth of genomic and phenotypic data available for nearly all known yeast species described in the subphylum Saccharomycotina ^51^ and cross-referencing with the ecological data available from the cactus-yeast-*Drosophila* system ^35–37,52–60^, we employed a high throughput framework to detect signatures of convergent evolution in 17 independently evolved lineages of cactophilic yeasts. Using a machine learning algorithm, we uncovered distinctive phenotypic traits enriched among cacti-associated yeasts, including the ability to grow at high (≥ 37°C) temperatures. We found that thermotolerance might be related to duplication and distinctive rates of evolution in genes impacting the integrity of the cell envelope, some of which are under positive selection in distantly related cactophilic clades. Gene family evolution analyses identified gene duplications and HGT events involving plant cell wall-degrading enzymes in distinct clades, suggesting adaptations associated with feeding on plant material. These results reveal that convergence to cactophily by distinct lineages of Saccharomycotina yeasts was accomplished through diverse evolutionary mechanisms acting on distinct genes, some of which might be associated with similar biological functions. Interestingly, we found that several cacti-associated yeasts and close relatives are emerging opportunistic human pathogens, raising the hypothesis that fungi inhabiting certain wild environments are preadapted for opportunistic pathogenicity. More broadly, we advocate for a methodological framework that couples diverse lines of genomic, phenotypic, and ecological data with multiple analytical approaches to investigate the plurality of evolutionary mechanisms underlying ecological adaptation.

## Results

### Yeast cactophily likely evolved independently 17 times

We examined the ecological association of yeasts with the cacti environment across a dataset of 1,154 strains from 1,049 yeast species ^51,61^. Yeast-cacti associations vary substantially in their strengths ^35^. Some cacti-associated yeast species are more cosmopolitan, being commonly isolated from cacti but also other environments (henceforth referred to as transient), whereas others are strictly cactophilic, defined as those almost exclusively isolated from cacti (Table S1; note that this classification is based on the available ecological information, which may be impacted by sampling bias and other sampling issues – it is possible that strictly cactophilic species could also be found in other, yet unsampled, environments). We observed that strictly cactophilic species are found across almost the entire Saccharomycotina subphylum spanning from the Trigonopsidales (i.e., *Tortispora* spp.) ^33^ to the Saccharomycetales (i.e., *Kluyveromyces starmeri*) ^52^ (Fig. 1). Using these species in an ancestral state reconstruction, we inferred a total of 17 origins for the evolution of cactophily (Fig. S1).

**Fig. 1.**
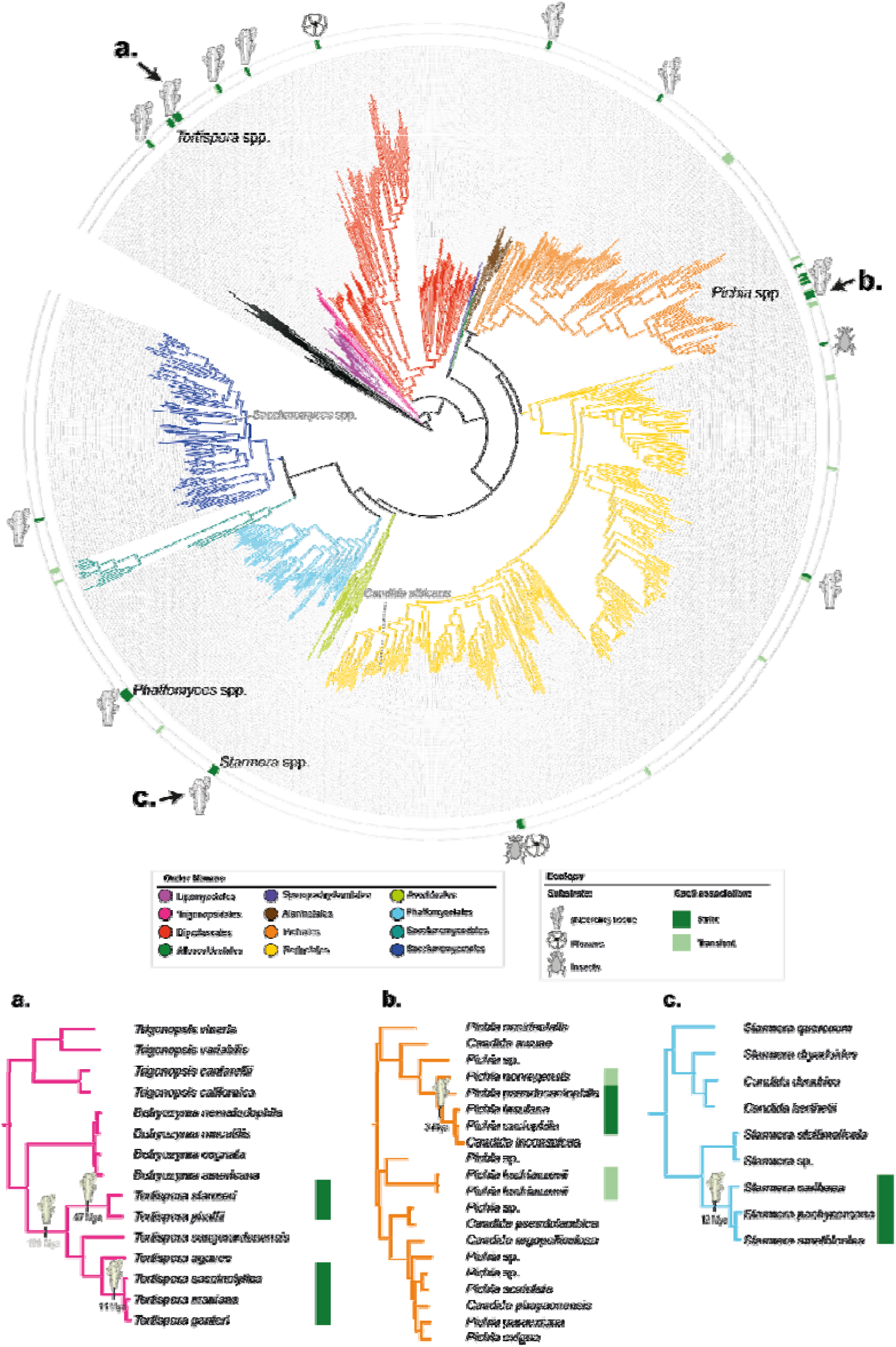
Yeast cactophily originated repeatedly and at different times. (top) Genome-wide-based phylogeny of the subphylum Saccharomycotina ^51^ depicting the different types of ecological association of strictly cactophilic yeast species with the cacti environment: (necrotic) cacti tissues, which include both *Opuntia* spp. and columnar cacti, cacti flowers, and cacti-visiting insects. Cactophilic lineages and well-known yeasts, such as *Saccharomyces* spp. and *Candida albicans*, are noted on the phylogeny. **a., b.,** and **c.**. Subtrees of three cactophilic clades: **a.** *Tortispora*, **b.** *Starmera*, and **c.** *Pichia*. Estimated times of origin, as determined in ^51^, for the emergence of cactophily are represented for these three example clades.

Cactophily is found in single species belonging to different orders, but it also involves nearly entire genus (i.e., *Tortispora*). Specifically, seven of the 17 instances of cactophily evolution involve clades containing two or more species while the remaining 10 involve single species, suggesting that different taxa evolved this ecological association at different times (Fig. 1). For instance, cross referencing relaxed molecular clock analyses of the yeast phylogeny ^51^ with ancestral state reconstructions suggests that cactophily in *Tortispora* likely emerged twice, once in the most recent common ancestor (MRCA) of *T. starmeri/T. phaffi* around 47 Mya and in the MRCA of *T. caseinolytica/T. mauiana/T. ganteri* around 11 Mya (Fig. 1). An alternative hypothesis would place the emergence of cactophily in the MRCA of the genus around 180 Mya, which is inconsistent with the 35 Mya estimated origin of the Cactaceae family ^62^. Cactophily in the genus *Starmera* and the *Pichia cactophila* group emerged more recently, most likely around 12 and 3 Mya, respectively (Fig. 1).

Cacti-associated yeasts seemingly exhibit significant niche partitioning (Table S1). For example, *T. ganteri* is typically isolated from columnar cacti, while its close relative *T. caseinolytica* is more commonly found in *Opuntia* spp. ^33^. Furthermore, *Pichia cactophila* is considered a generalist cactophilic yeast, being widely distributed across a wide range of cacti species ^37^, while closely related *P. heedii* has been predominantly found in association with certain species of columnar cacti ^39^. However, many species (e.g., *T. starmeri* or *P. insulana*) alternate between the two types of cacti ^33,58^, similar to some *Drosophila* species ^41^. Other species, such as *Wickerhamiella cacticola* or *Kodamaea nitidulidarum*, are associated with cacti flowers and/or flower-visiting insects, like beetles, and not with necrotic cacti tissues ^59,60,63^.

### Detecting signatures of convergent evolution in cactophilic yeasts

We envision three distinct scenarios that may capture how different yeast lineages convergently evolved cactophily (Fig. 2A):

**Fig. 2.**
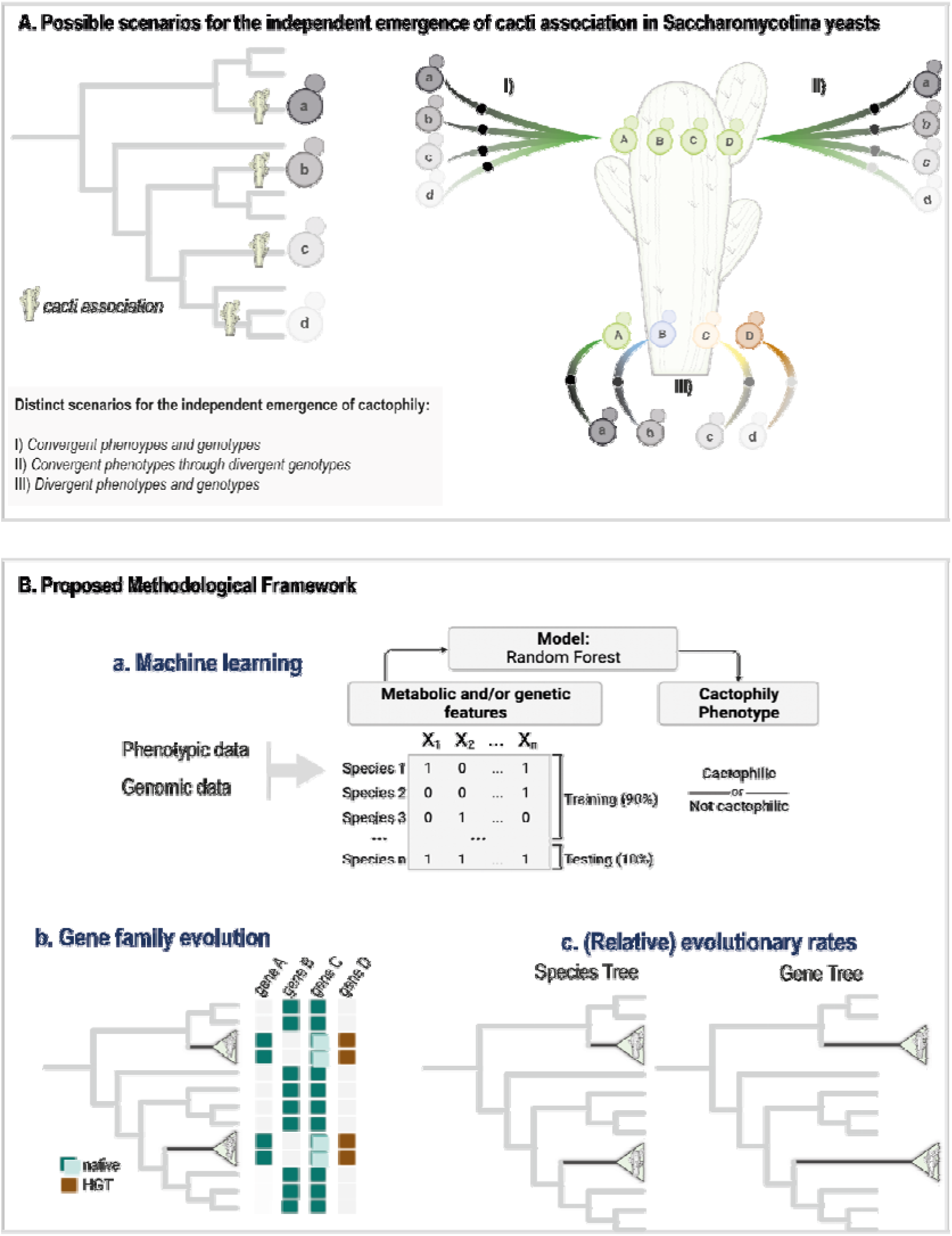
Alternative scenarios for the evolution of yeast cactophily and methodological framework employed in this study. (A) Methodological framework for investigating signatures of convergent evolution of yeast cactophily (B). A) Association with cacti evolved multiple times in yeasts from distinct genetic backgrounds, which are represented by distinct shades of grey. Convergent adaptation to the cacti environment might have involved convergence at both phenotypic and genomic levels (Scenario I), convergence at the phenotypic level but through distinct molecular paths (Scenario II) or involved lack of convergence at both phenotypic and genotypic levels (Scenario III). Each of these scenarios is tested using a methodological framework (B) involving (a) machine learning approach wherein a model is trained to distinguish cactophilic from non-cactophilic yeasts from genomic and phenotypic features; (b) gene family evolution to find evidence of gene gains, duplications, losses, and HGT that might have occurred in cactophilic yeasts; (c) evaluation of signatures of positive selection (omega, _ω_) and of changes in relative evolutionary rates in branches leading to cactophilic clades. This strategy will expose both genomic and phenotypic traits associated with cactophily (a, b) and highlight genes that are evolving slower or faster than the average of the genome or are under positive selection in cactophilic vs. non-cactophilic species (c).

#### Scenario I: Convergent phenotypes and genotypes

Selective pressures associated with the cacti environment (e.g., high temperature, desiccation, or presence of toxic compounds) favor similar phenotypic traits that evolved through the same genomic mechanisms;

#### Scenario II: Convergent phenotypes through divergent genotypes

Selective pressures associated with the cacti environment favor similar phenotypic traits across cactophilic species, but different evolutionary mechanisms (e.g., gene duplication, HGT) contribute to phenotypic convergence of different lineages. In this scenario, similar phenotypes emerge through distinct evolutionary trajectories;

#### Scenario III: Divergent phenotypes and genotypes

Distinct phenotypic landscapes are explored by distinct clades when thriving in the same environment (niche partitioning); different evolutionary mechanisms contribute to these phenotypes.

To explore which scenario(s) best reflect(s) the process of yeast adaptation to the cacti environment, we developed a framework for identifying signatures of adaptation and convergence from high-throughput genomic and phenotypic data ^51^ (Fig. 2B). Specifically, we performed: machine learning to identify phenotypic and genetic commonalities that distinguish cactophilic from non-cactophilic yeasts; genome-wide family analyses to identify patterns of gene presence/absence due to gene duplication, gene loss, and HGT; and genome-wide evolutionary rate analyses to detect signatures of convergence in evolutionary rates and of positive selection in individual genes (Fig. 2B), which have been also frequently implicated in adaptive evolution ^5,19,21,22,64^. We applied this methodological framework to study convergence in ecological specialization in yeasts but note that it can be applied more generally to study the process of convergent or adaptive evolution.

### Specific metabolic and genomic traits predict cactophily

To investigate if cactophilic yeasts share similar phenotypic and/or genomic traits, we used a dataset of 1,154 yeast strains ^51^, from which 52 are either strictly cactophilic (rarely found in other environments, n=31) or transient (frequently isolated from cacti but cosmopolitan, n=21) (Table S1). Functional genomic annotations (KEGG – 5,043 features) and physiological data (122 features) were retrieved ^51^ and used as features in a supervised random forest (RF) classifier trained to distinguish cactophilic from non-cactophilic yeasts. By training 20 independent RF runs using randomly selected balanced datasets (52 non-cactophilic species randomized each time and the 52 cactophilic species), we correctly identified an average of 38 cactophilic species (Fig. 3A), yielding an overall accuracy and precision of 72% and 73%, respectively. Species incorrectly assigned in 10 or more independent runs were equally distributed across strictly and transiently cactophilic groups (seven in each) (Table S2). We next repeated the analysis considering only strictly cactophilic species (transient were considered non-cactophilic) and obtained slightly higher accuracy (76%) and precision (76%) (Fig. 3B). Notably, correct classifications were obtained across phylogenetically distantly related genera (e.g., *Tortispora*, *Phaffomyces* or *Pichia*) (Fig. 3A), while incorrect classifications were obtained for species belonging to cactophilic clades in which correct classifications were obtained; for instance, despite being closely related, we obtained both correct and incorrect classifications for cactophilic species belonging to the *Phaffomyces* genus (highlighted in light blue, Fig. 3A). These results suggest that the phylogenetic relatedness between some cactophilic species did not interfere with the accuracy of the RF classifier.

**Fig. 3.**
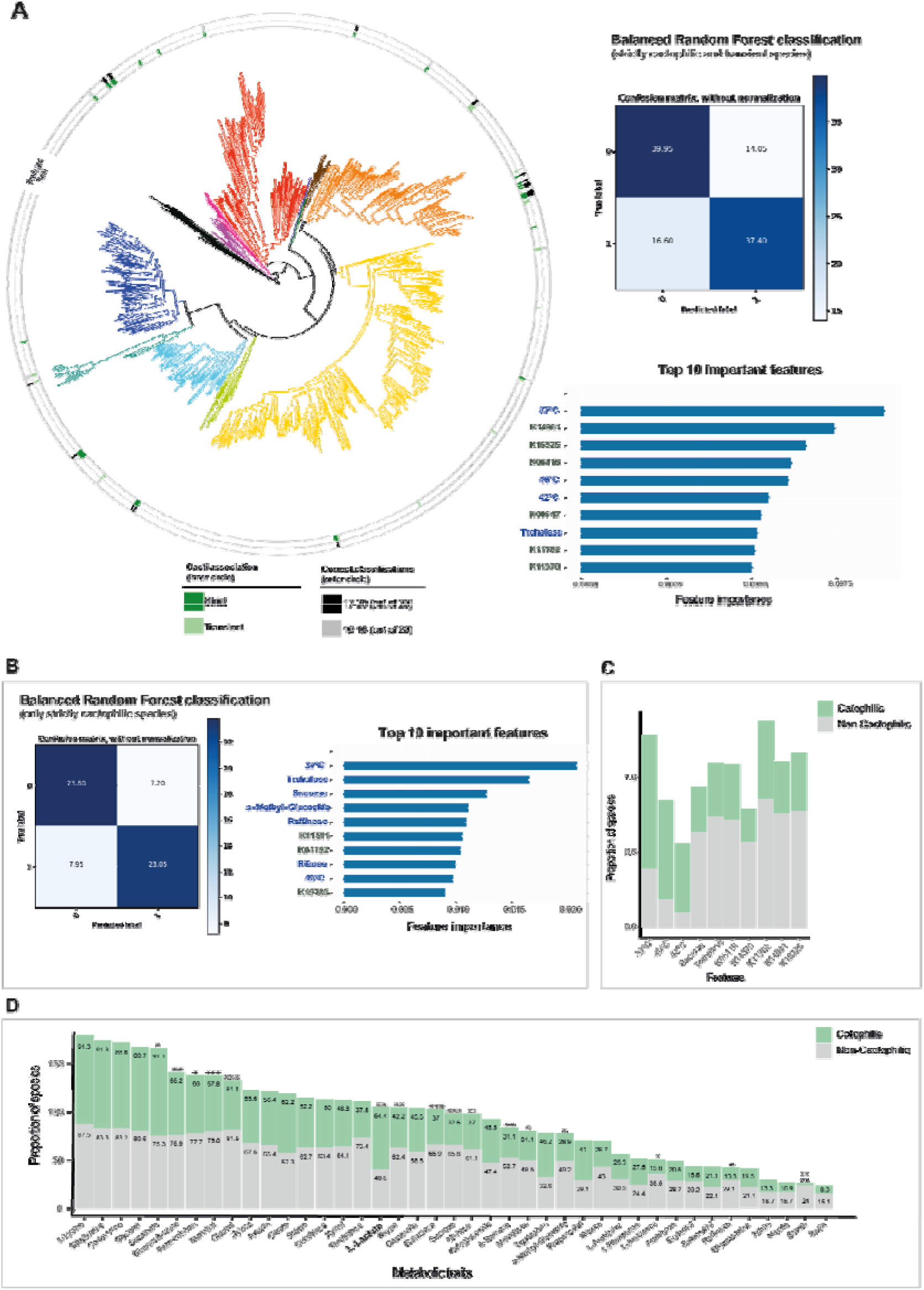
Cactophilic and non-cactophilic yeasts can be predicted from genomic and metabolic data with good accuracy. A) (left) Distribution of correct random forest (RF) classifications of cactophilic yeasts across the Saccharomycotina phylogeny. (right) Confusion matrix showing the average number of true positives (37.40), true negatives (39.95), false positives (16.60), and false negatives (14.05) across strictly cactophilic and transient species resulting from 20 independent RF runs. On the bottom, the top ten most important metabolic (presence or absence of growth; in blue) and genomic (presence or absence of KEGG; in grey) features for the RF classifier ranked according to their importance scores are shown. B) Confusion matrix and top ten most important features for the RF classifier using only strictly cactophilic species. C) Distribution of the proportion of the most important features to predict cactophily (strict cactophilic and transient) and non-cactophilic species for the entire dataset of 1,154 yeasts. D) Distribution of the proportion of metabolic traits (representing presence or absence of growth under the conditions shown) ^51^ across cactophilic (strict and transient) and non-cactophilic species. Only metabolic traits for which no more than 50% of data were missing were considered. Additionally, only metabolic traits exhibiting more than 10% prevalence in one of the groups (cactophilic or non-cactophilic) are shown. Statistically significant differences (Chi-Squared Test; * p-value < 0.05, ** p-value < 0.01, *** p-value < 0.001) between the proportions of cactophilic and non-cactophilic species able to grow in each carbon/nitrogen source are shown.

We next examined the top features that significantly contributed to the RF classifier. The feature with the highest relative importance was growth at 37°C either when analysing all cactophilic species (Fig. 3A) or just strictly cactophilic species (Fig. 3B). Ability to grow at high temperatures was previously found to be prevalent in cacti-associated yeasts ^38,65^ and is also an adaptive feature of cactophilic *Drosophila* ^48,66^. Growth at 40°C and 42°C were also among the top ten most important features, supporting the hypothesis that thermotolerance is a distinctive feature of cactophilic species. In fact, 90%, 66%, and 46% of cactophilic yeasts can grow at 37°C, 40°C and 42°C, respectively, compared to only 39%, 19% and 10% of non-cactophilic yeasts (Chi-Squared Test, p < 0.01) (Fig. 3C).

Analysing the top 100 most important metabolic features, we observed that 74 are less common in cactophilic species compared to non-cactophilic species (Table S2). For instance, trehalose assimilation is more rarely found in cactophilic species (∼25%) than non-cactophilic species (∼74%) (Chi-Squared Test, p-value < 0.01) (Fig. 3C). Trehalose generally accumulates during numerous stress conditions including heat stress ^67,68^. When cells return to a more favourable condition, the accumulated trehalose is hydrolysed into glucose by the trehalase Nth1. Inactivation of *NTH1* by mutations, and therefore subsequent impairment of trehalose hydrolysis, can be one of the outcomes of the heat stress response in experimentally evolved strains of *S. cerevisiae* under high temperature stress ^69,70^, suggesting that deficient trehalose hydrolysis can be beneficial under long-term thermal stress conditions. However, *NTH1* is generally present in cactophilic genomes, suggesting that absence of *NTH1* does not explain impairment in trehalose assimilation in these species. Other top important metabolic features, such as sucrose assimilation, are also more rarely found in cactophilic species (Chi-Squared Test, p-value < 0.001) (Table S2, Fig. 3C), echoing the general trend of a narrower spectrum of carbon sources assimilated by cactophilic species compared to their non-cactophilic counterparts (Fig. 3D). However, some metabolic traits are more frequently found among cacti-associated yeasts, such as assimilation of lactate (Chi-Squared Test, p-value < 0.01) (Fig. 3D), which was previously found to be positively associated with the cacti environment ^38,65^.

Among the most important genomic features were presence or absence of genes involved in multiple distinct functions: K15325 (splicing), K06116 (glycerol metabolism), K01192 (N-glycan metabolism), K00547 (amino acid metabolism), K11762 (chromatin remodelling), K11370 (DNA repair), K11511 (DNA repair), or K19783 (post-replication repair). All these features are less common in cactophilic than in non-cactophilic species (Table S2). One interesting exception is K03686, a Hsp40 family protein encoded by 85% of cactophilic and 57% of non-cactophilic species (Table S2).

### HGT and duplication of cell wall degrading enzymes in cactophilic yeasts

We next looked for genes that might be implicated in cactophily by examining gene family evolution ^71^ across three groups that contained two or more cactophilic lineages. We constructed three distinct datasets (Table S3) containing species of interest and closest relatives within the Lipomycetales/Dipodascales/Trigonopsidales orders (referred to as LDT group, including *Tortispora* spp., *Dipodascus australiensis*, *Magnusiomyces starmeri*, *Myxozyma mucilagina*, and *Myxozyma neglecta*), Phaffomycetales (including *Starmera* spp. and *Phaffomyces* spp.) and Pichiales (including two distinct *Pichia* spp. cactophilic clades).

We focused on gene duplications and gene gains (e.g., HGT), as gene losses are usually not reliably estimated due to annotation and sampling issues or inaccurate gene family clustering ^71^. Using gene tree – species tree reconciliation analyses implemented in GeneRax ^71^ we first examined genes with evidence of duplication in at least one cactophilic species belonging to each group. We cross-referenced the candidate genes with their respective functional annotations (KEGG) ^51^ and found 11 KEGG families with evidence of duplication in all three groups and 29–37 families with evidence of duplication in two of the three groups (Table S4). Among these gene families, we found duplicated cellulase-encoding genes (K01210) in *Dipodascus australiensis* (LDT group) and *Starmera* species (Phaffomycetales). For the latter species, however, we could not recover complete gene sequences as they were located either at the end of scaffolds or in short scaffolds. Nevertheless, BLASTp searches against the non-redundant (nr) NCBI database indicate that these sequences were endoglucanase-like enzymes similar to those found in filamentous fungi (e.g., KOS23166.1) and are rarely found in yeasts^72^.

Duplication of another gene involved in plant cell wall degradation, encoding a rhamnogalacturonan endolyase (K18195), was detected in the cactophilic *P. antillensis*, *P. opuntiae*, and *Candida coquimbonensis* (Phaffomycetales, Table S4). These species contained two copies of this gene compared to their closest relatives, which contained only one (Fig. 4A). These enzymes are responsible for the extracellular cleavage of pectin ^73^, which along with cellulose, is one of the major components of plant cell wall. Pectin lyase ^74^ activities have been rarely reported among yeasts; therefore, we assessed the distribution of the rhamnogalacturonan endolyase across the 1,154 proteomes using a BLASTp search (e-value cutoff e^-3^) and found that it displayed a patchy distribution, being found in fewer than 60 species (Fig. 4A).

**Fig. 4.**
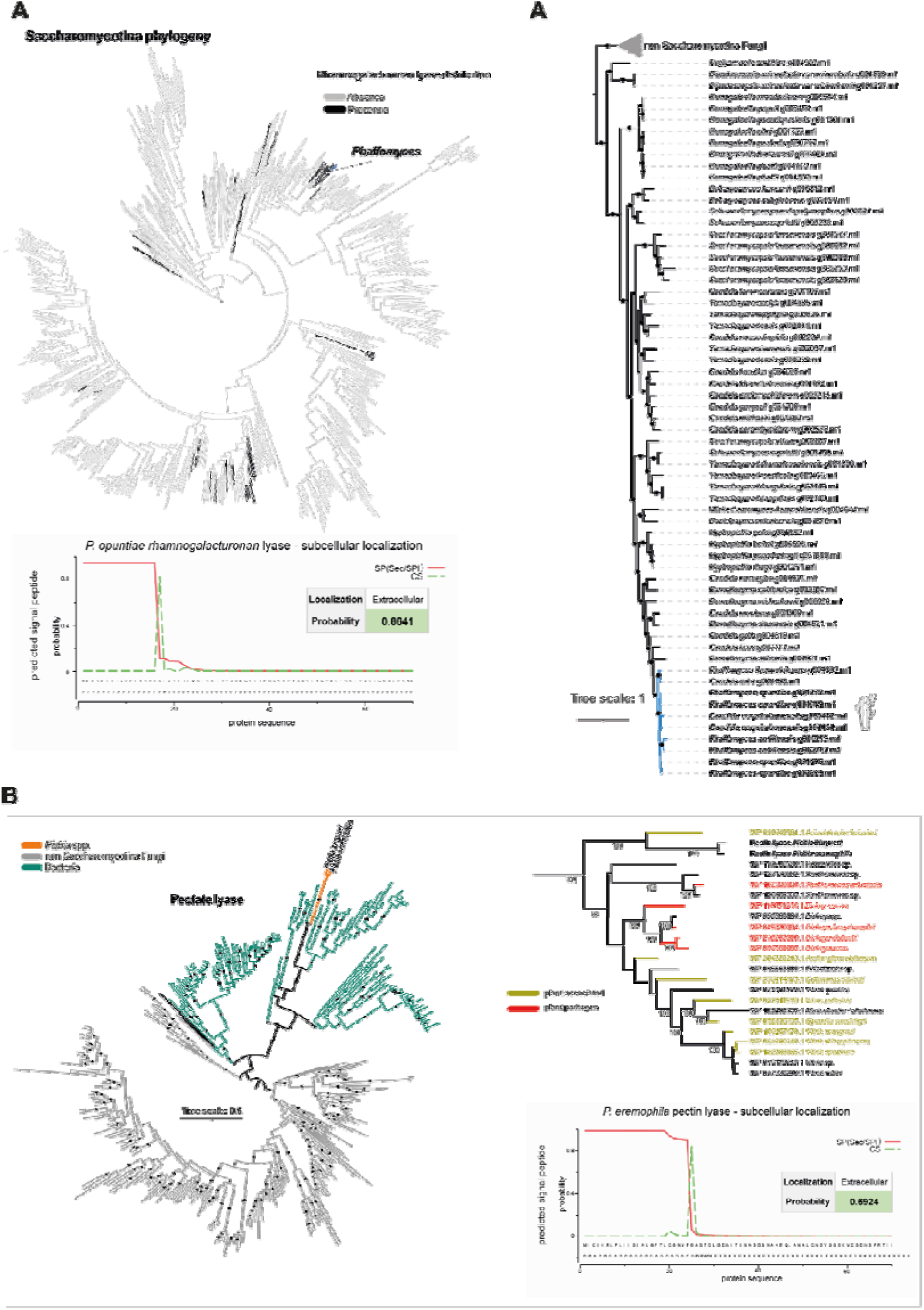
Duplication and horizontal gene transfer of plant cell wall-degrading enzymes in cactophilic species. A) Distribution of rhamnogalacturonan endolyases across the 1,154 yeast genomes (presence in black and absence in grey). (on the right) Phylogenetic tree of yeast rhamnogalacturonan endolyase and closest relatives, highlighting the duplication events in cactophilic *Phaffomyces* species. Phylogeny was constructed in IQ-TREE v2.0.6 (-m TEST, -bb 1,000) ^81,82^. Branch support (bootstrap >= 90) is represented as black circles. A BLASTp search against the NCBI nr database (selecting 500 hits) was performed using *P. opuntiae* g001049.m1 protein sequence as query. BLASTp was also performed against the yeast dataset of 1,154 proteomes and all significant hits were retrieved (e-value cutoff e^-3^). In *P. opuntiae*, there are two additional partial sequences (g001496.m1 and g001773.m1, 163 amino acids) that only partially overlap (from 39 to 163 overlapping amino acids) with the remaining nearly complete sequences (g002229.m1: 468 amino acids and g001049.m1: 610 amino acids). Prediction of subcellular localization according to SignalP and Deeploc ^72,83^ is shown in the panel below. B) Phylogenetic tree of closest related sequences to pectin lyases from *P. eremophila* and *P. kluyveri.* (on the right) Pruned tree highlighting the ecological association of bacteria species harboring the closest related pectin lyase sequences to *P. eremophila/P.kluyveri* proteins. Prediction of subcellular localization^72,83^ is shown in the panel below.

Interestingly, by inspecting orthogroups that uniquely contained cactophilic species, we found a gene encoding a pectate/pectin lyase was uniquely found in *P. eremophila* (strictly cactophilic) and *P. kluyveri* (transient), which are commonly isolated from rotting cacti tissues. Sequence similarity searches across the entire dataset of 1,154 yeast genomes confirmed that this gene is absent from all other species. Pectate lyases are also extracellular enzymes involved in pectin hydrolysis and plant cell wall degradation. Consistent with this function, these enzymes are mostly found among plant pathogens and plant-associated fungi and bacteria ^75,76^, and their activity has only been reported in a handful of Saccharomycotina species ^74^. Phylogenetic analyses showed that the two yeast sequences are nested deeply within a clade of bacterial pectin lyases (Fig. 4B). The most closely related sequence belongs to *Acinetobacter boissieri* ^77^, which has been frequently isolated from plants and flowers, and to *Xanthomonas* and *Dickeya*, two genera of plant pathogenic bacteria ^78,79^.

Consistent with their function, HGT-derived pectin lyases as well as cellulases and rhamnogalacturonan endolyases, were predicted to localize to the extracellular space based on primary sequence analyses ^72^ (Fig. S2, Fig. 4). Pectin lyase enzymatic activity was previously detected in *P. kluyveri* strains associated with coffee fermentation ^76,80^, suggesting that the identified HGT-derived pectin lyase is likely responsible for this activity.

### Convergent accelerated rates in heat resistance-related genes

We next specifically looked for evidence of convergent evolutionary rates ^84^, another indicator of adaptation ^21,22^. For this analysis, we selected all cactophilic species found within the Pichiales, Phaffomycetales and the LDT group.

Correlation analyses between relative evolutionary rates (RER) and cactophily across ancestral (anc) and terminal branches revealed changes in evolutionary rate associated with the evolution of cactophily. Specifically, we inferred that 20 / 3,029 gene families in the LDT group are under accelerated evolution (evolving significantly faster than the average gene) and 33 / 3,029 have undergone decelerated evolution (evolving significantly slower than the average gene) (Table S5, Fig 5A). In the Pichiales, 14 / 2,204 showed evidence of acceleration and 25 / 2,204 of deceleration (Fig. 5A). In the Phaffomycetales, we found 32 / 3,550 accelerated genes and 30 / 3,550 decelerated genes (Table S5, Fig. 5A).

**Fig. 5.**
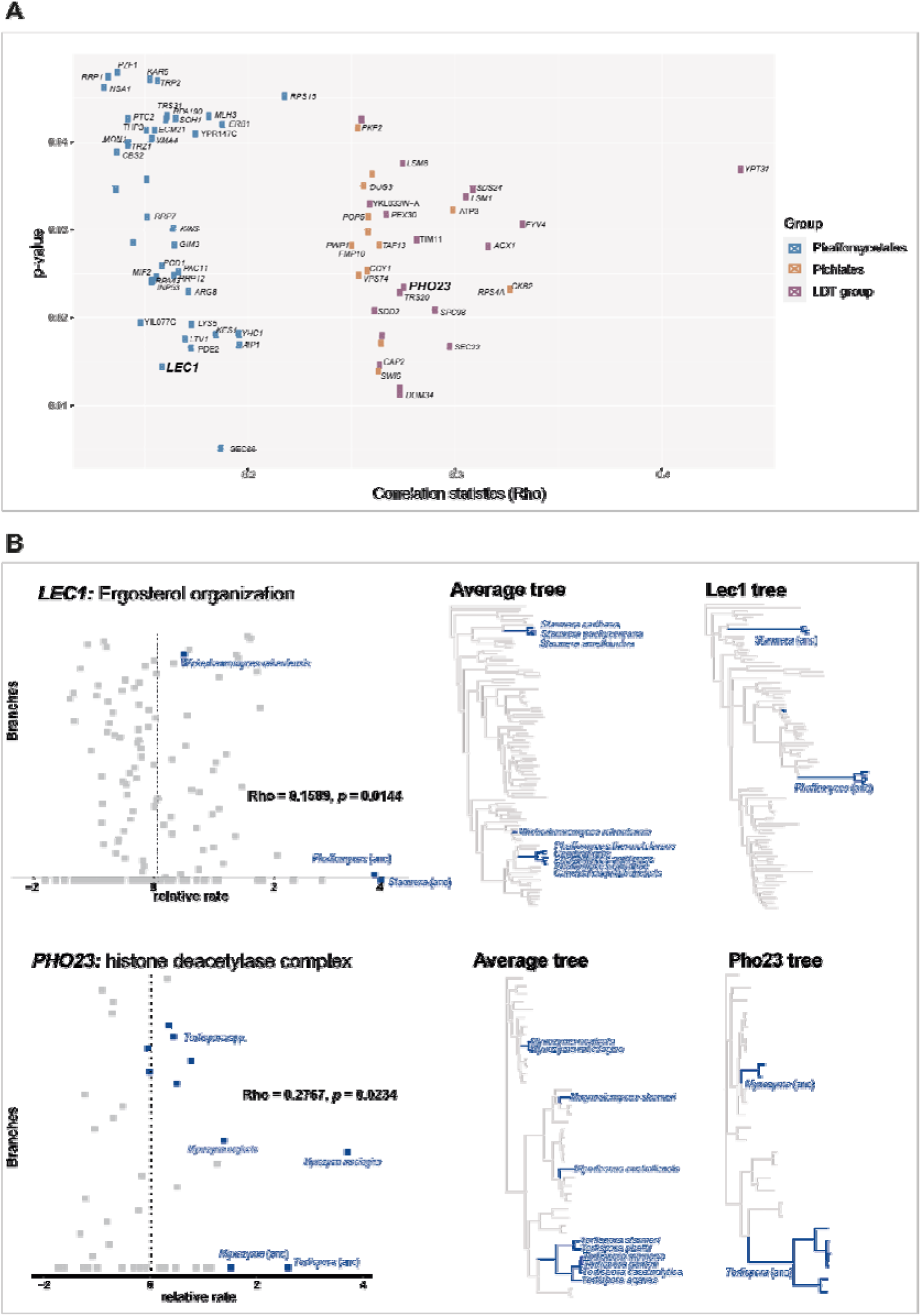
Genes involved in maintenance of the cell envelope show accelerated evolution in cactophilic clades. A) Genes with accelerated evolutionary rates in the three groups of cactophilic species inspected. Correlation statistics (Rho) for converge between accelerated rates of evolution in specific genes and cactophilic as well as the respective statistical significance of the correlation (p-value) are displayed. B) Relative evolutionary rates (RER) and respective fixed-topology phylogenies for genes related with integrity of the cell membrane (*LEC1*) and heat stress (*PHO23*).

The accelerated genes are associated with varied cellular functions. In Pichiales, 7 out of 14 accelerated genes impact heat resistance according to large-scale studies ^85^. For instance, *SWI6*, encodes a transcription factor that induces transcription during heat stress, and deletion of this gene causes several impairments in the resistance to multiple stresses, including heat ^86^ and cold ^87^. Among the accelerated genes in Phaffomycetales, we found *LEC1*, which was recently associated with ergosterol organization (Fig. 5B) ^88^.

Inspecting the literature for phenotypes associated with either null or conditional mutants ^85^, we found that, irrespective of their function, 12 / 20 genes that exhibited accelerated rates in the LDT group are involved in either heat and/or desiccation resistance. For instance, loss of function mutations in *CAP2*, which encodes part of a capping complex involved in barbed-end actin filament capping and filamentous growth, are associated with heat sensitivity and abnormal chitin localization leading to aberrant cell morphology ^89,90^. Another gene identified as having accelerated evolutionary rates in the LDT group was *PHO23* (Fig. 5B), which was found to be required for the growth of *S. cerevisiae* during heat shock ^86^. Genes that underwent decelerated evolution also play multiple roles (Table S5), and some are involved in essential functions such as DNA repair, cell cycle and splicing or encoding ribosomal subunits.

We next assessed the occurrence of positive selection across cactophilic clades using branch-site tests of rates of nonsynonymous (dN) and synonymous substitutions (dS). These tests were performed separately for each of the five cactophilic subclades within the major lineages selected: *Pichia* A and *Pichia* B clades (Pichiales), *Starmera* and *Phaffomyces* clades (Phaffomycetales), and *Tortispora* clade (Trigonopsidales) (Fig. S3). Only genes for which no evidence of positive selection was found in the non-cactophilic sister clades (p-value < 0.05) were considered (Fig. S3). With this conservative approach, we found evidence for positive selection for 292 / 2,384 genes examined in *Tortispora*; 328 / 2,175 in *Pichia* A; 259 / 2,155 in *Pichia* B; 90 / 1,685 in *Phaffomyces*; and 105/1,602 in *Starmera* (Table S6). Importantly, signatures of selection (significantly higher ω values than 1 in branches leading to cactophilic clades) can stem from either high dN or low dS values ^91^, and we did not specifically distinguish between the two scenarios. Genes under positive selection showed limited overlap; six genes were under positive selection in three clades and 134 genes in two clades, while no genes presented evidence for positive selection in all five clades.

While the distribution of presence / absence of the trehalase gene *NTH1* was not associated with the low prevalence of trehalose assimilation in cactophilic species (Fig. 3C), *NTH1* was among the genes under positive selection in the two *Pichia* clades (Table S6). The 140 genes with evidence of positive selection in two or three clades were enriched in multiple biological functions including chemotropism, translation and splicing, carnitine metabolism, lactate oxidation, and ergosterol biosynthesis (Table S7). More specifically, 6/140 genes were involved in several steps of ergosterol biosynthesis (*ERG1*, *ERG8*, *ERG24*, *ERG26*, *UPC2, SIP3*) (Fig. 6). *ERG13* was also under positive selection in the *Starmera* clade. Ergosterol is involved in stabilising cell membranes during heat stress and therefore has a major role in the tolerance to numerous stresses in fungi ^68,92–94^.

**Fig. 6.**
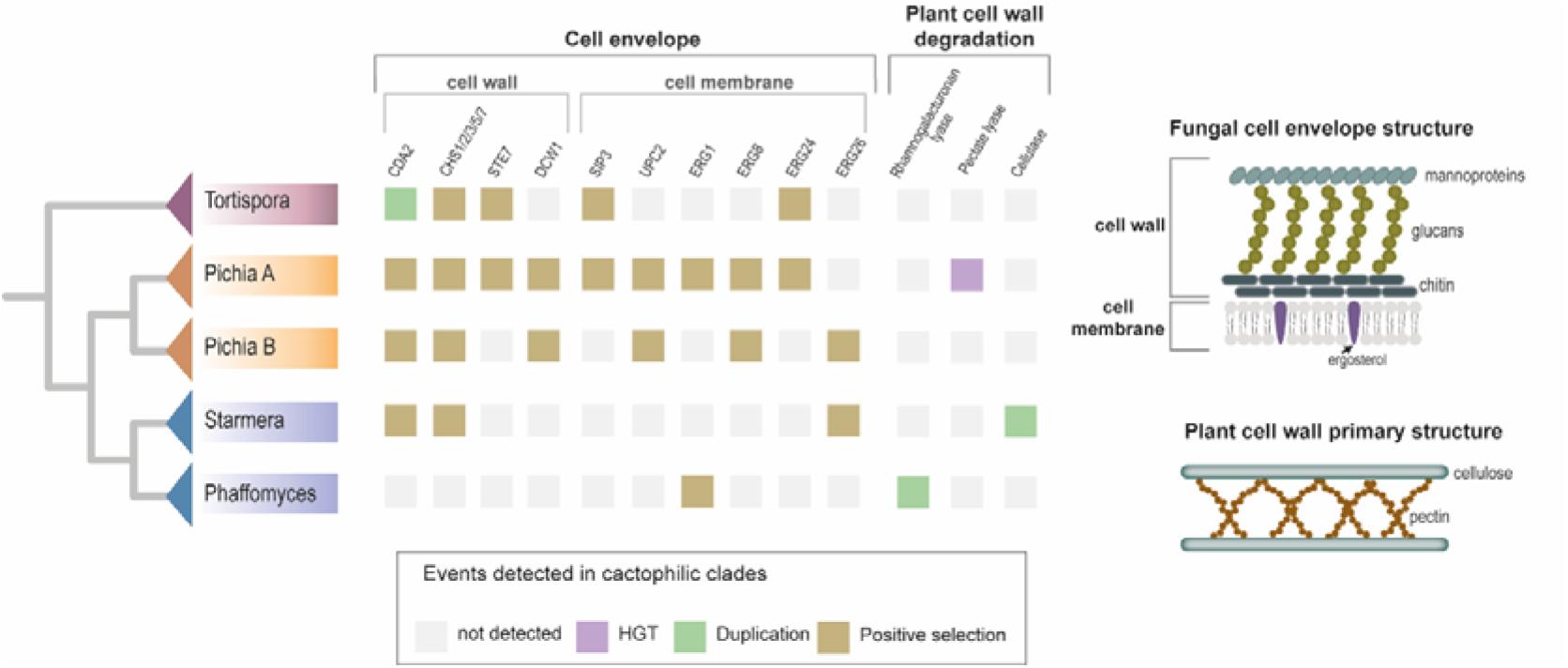
Signatures of convergent molecular evolution across cactophilic clades. Genes and functions for which detection of distinctive evolutionary alterations (positive selection, duplication, and HGT) in more than one cactophilic clade are highlighted. The distinct evolutionary events are represented by squares filled with different colors, as indicated in the key. Schematic representations of the fungal cell envelope and plant cell wall are shown as many of the genes with signatures of convergence are associated with functions impacting these two structures.

Other genes were specifically involved in cell wall biosynthesis and integrity (Table S6). For instance, *CDA2*, which encodes a chitin deacetylase involved in the function of the fungal cell wall ^95,96^, was under positive selection in the stem branches of three distinct cactophilic clades (Fig. 6) and was duplicated in the *Tortispora* clade (Fig. 6).

Other cell wall-related functions were found among the group of genes that showed evidence for positive selection in two or more clades, namely chitin synthases *CHS1*, *CHS2*, and *CHS3* and chitin-related genes *CHS5* (involved in Chs3 transport from the Golgi to the plasma membrane) and *CSH7* (involved the export of Chs3 from the endoplasmic reticulum) and *UTR2* (chitin transglycosylase). *DCW1* (encoding a mannosidase required for cell wall formation and resistance to high temperatures) ^97^, *STE7* (encoding a signal transducing MAP kinase involved in cell wall integrity and pseudohyphal growth) ^98^, and *AYR1* (encoding a bifunctional triacylglycerol lipase involved in cell wall biosynthesis) ^99^, were also found among the genes under positive selection in two cactophilic clades. Mutations in these genes have been associated with cell wall defects and increased heat sensitivity in *S. cerevisiae* ^100–103^ and differential transcriptional responses to heat stress have also been documented for some of these genes and functions ^104^.

### Some genes under positive selection in cactophilic clades show evidence of codon optimization

To infer the transcriptional activity of cactophilic yeasts, we determined gene-wise relative synonymous codon usage (gw-RSCU), a metric which measures biases in codon usage that have been shown to be associated with expression level ^105^, and examined the top-ranked genes (95^th^ percentile) in cactophilic species (Table S8). Top-ranked genes include many encoding ribosomal subunits and histones, which are known to be highly expressed and codon-optimized in *S. cerevisiae* ^106^. We found that the chitin deacetylase gene *CDA2* and genes involved in ergosterol biosynthesis (namely *ERG2*, *ERG5*, *ERG6*, and *ERG11*) were among the genes that fell within the 95^th^ percentile rank for gw-RSCU in multiple cactophilic species. To ascertain whether these genes also show signatures of codon optimization in closely related non-cactophilic species, we determined their respective gw-RSCU percentile ranks. While no clear pattern was observed for *ERG* genes (these genes were also highly ranked for gw-RSCU in non-cactophilic species), we observed that *CDA2* is particularly highly ranked in *Phaffomyces*, *Starmera*, and *Pichia* clades compared to their closest relative non-cactophilic species (Fig. S4). *CDA2* showed evidence for positive selection in both *Pichia* clades and *Starmera*, suggesting that distinctive synonymous and/or nonsynonymous might have resulted from translational selection for optimized codons due to higher gene expression.

### Cactophily as a launching pad for the emergence of opportunistic human pathogens?

Thermotolerance is a key shared trait by human fungal pathogens ^27,107–109^. Interestingly, several cactophilic or closely related species are emerging human opportunistic pathogens (Fig. 7, Fig. S5). Examples include *Candida inconspicua* and *Pichia norvegensis* ^110^, which cluster within the *Pichia cactophila* clade (Fig. 7), and *Pichia cactophila*, which was also isolated from human tissue ^111^. Cases of fungemia have also been associated with *Pichia kluyveri* ^112^, a transient species belonging to a separate clade within the Pichiales. *Kodamaea ohmeri*, which has also been isolated from the cacti environment ^52^ and is closely related to the cactophilic *Ko*. *nitidulidarum* and *Ko. restingae*, is also an emerging human pathogen with a significant mortality rate ^113–115^.

**Fig. 7.**
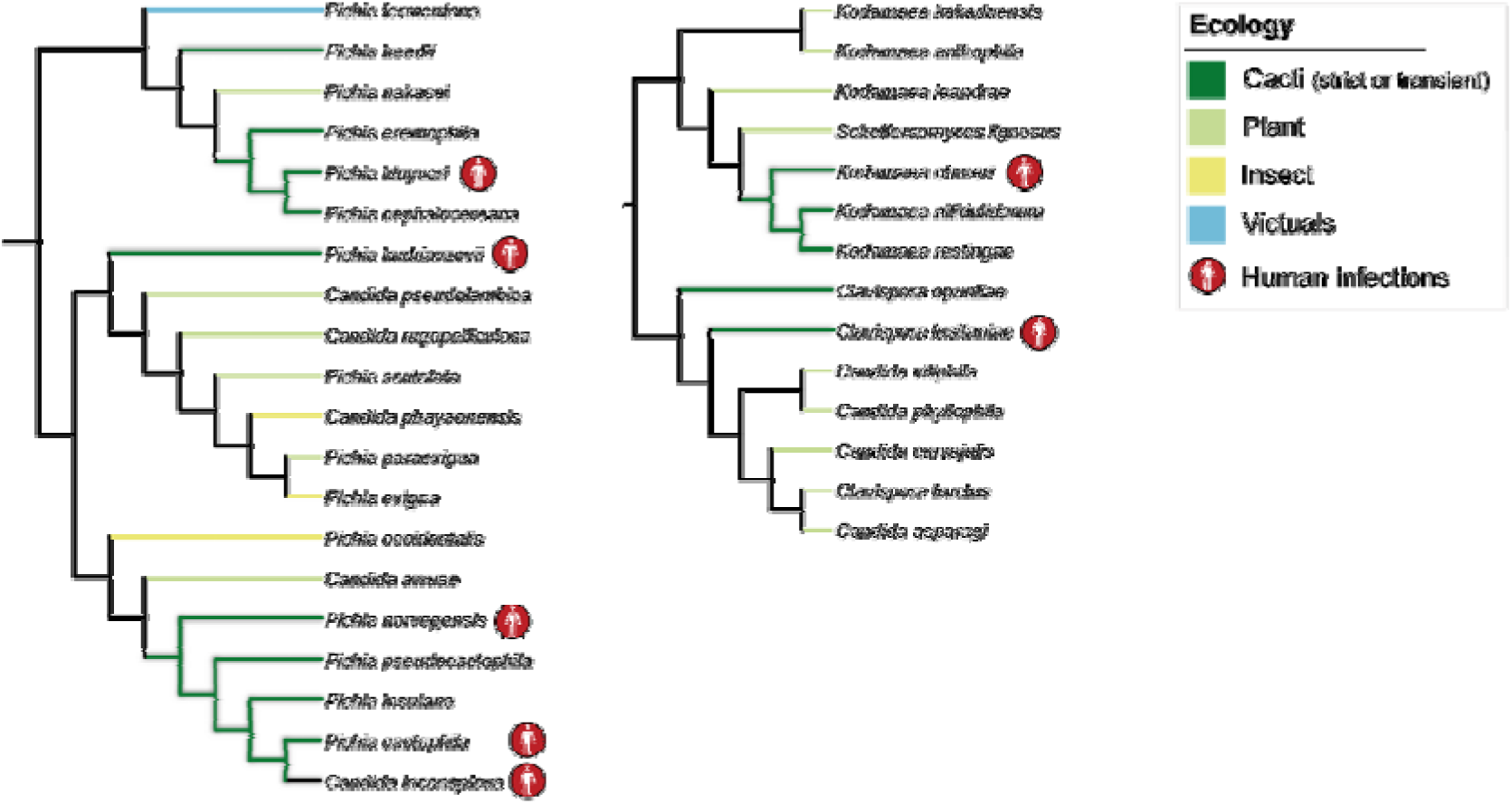
Example cactophilic lineages that contain human opportunistic pathogens. Ecological associations of cactophilic species and their closest relatives are represented, highlighting examples of species associated with human infections. Ecological information for additional cactophilic species and closest relatives is provided in Fig. S5.

In addition to thermotolerance, other aspects of the cactophilic lifestyle might be pre-adaptations for human pathogenicity. For instance, the sterol composition of the cell envelope has been implicated in fungal virulence ^94,116–118^. Mutations in genes involved in the ergosterol biosynthetic pathway are associated with antifungal resistance ^118–121^, making it a major target of antifungal drugs ^122^. We showed that the evolutionary rates of several genes involved in the ergosterol biosynthesis across multiple cactophilic clades have changed, suggesting that this pathway might be under a new selection regime in these lineages. However, the impact of these alterations remains to be elucidated.

## Discussion

By examining high-throughput genomic, phenotypic, and ecological data for 1,049 yeast species we unveiled multiple (∼17) independent occurrences of cacti association. The ability to grow at ≥ 37°C emerged as the strongest predictor of the cactophilic lifestyle. Being a polygenic trait, thermotolerance can arise through multiple distinct evolutionary trajectories ^123^. Heat stress generally affects protein folding and cell integrity and involves a complex response from multiple genes impacting the expression of heat shock proteins, the integrity of the cell wall and membranes, production of compatible solutes, repression of protein biosynthesis, and/or temporary interruption of the cell cycle ^67^. The expression of genes involved in cell wall biosynthesis and integrity is for instance affected when strains of *S. cerevisiae* are exposed to heat stress ^104^. Possibly in association with thermotolerance, we found genes involved in maintaining the cell envelope exhibiting evidence of positive selection, codon optimization, and duplication in multiple cactophilic clades.

We also found genomic fingerprints that indicate phenotypic convergence in the ability to feed on plant material. Acquisition and duplication of plant cell wall-degrading enzymes can be interpreted as adaptive and supports the involvement of cactophilic yeasts in the cacti necrosis process ^40^. Interestingly, the contrasting mechanisms employed, HGT of a bacterial pectin lyase and duplication of cellulases and a rhamnogalacturonan lyase, show that phenotypic convergence can arise through disparate molecular mechanisms. While we found possible cases of phenotypic and molecular convergence for thermotolerance (Scenario I, Fig. 2) and cases of possible phenotypic convergence through distinct molecular mechanisms in respect to plant cell wall degrading ability (Scenario II, Fig. 2), we found limited evidence for convergence at both phenotypic and genomic levels (Scenario III, Fig. 2). Based on these data, we conclude that cactophily can originate through multiple phenotypic and genetic changes, some commonly found and some more rare.

It was previously postulated that most cactophilic yeasts evolved from ancestors associated with plants ^37^ (Fig. S5), indicating that the ability to thrive in plant-related environments was already present in the ancestors of many of the lineages that evolved cactophilic lifestyle. However, cacti are native to arid and semiarid climates in the Americas ^124^, where extremely high (and low) temperatures and low humidity constitute crucial challenges. This reasoning aligns with our finding that thermotolerance is the phenotypic feature with the strongest signature of adaptation to cacti.

Compared to mammals, a lineage that has provided spectacular examples of parallel molecular evolution underpinning the independent emergence of convergent traits ^5,8,64,125–127^, yeasts exhibit far higher levels of genetic and physiological diversity ^26^. Consequently, the probability of finding overlapping evolutionary paths might be reduced as pleiotropic effects or mutational epistasis might be more prominent across divergent genetic backgrounds ^9,10^. It is also possible that collection of additional phenotypic data or evolutionary genomic analyses will revise our understanding of the nature of convergent evolution to cactophily. This caveat notwithstanding, these results present a first snapshot of the study of convergent evolution of an ecological trait in yeasts, employing multiple state-of-the-art methodologies that aim at looking into a wide range of evolutionary mechanisms, phenotypes, and genetic determinants. It also underlines the exceptional value of combining high throughput physiological, genomic, and ecological data to investigate still-pressing questions in evolutionary biology.

## Materials and Methods

### Species selection

Yeast species associated were selected from Opulente & LaBella et al. ^51^ by cross referencing ecological information available from the literature ^33,35–37,39,52,54,55,58–60,63,128,129^. Two different groups of cactophilic species were determined according to their degree of association with cacti: strictly cactophilic (species that are mainly isolated from cacti and very rarely isolated from other environments) or transiently cactophilic (species frequently found in cacti, but also frequently found in other environments or for which strong association with cacti was not clear from the literature) (Table S1). Species very rarely isolated from this environment were not considered as they could either represent misidentifications or have originated from stochastic events.

### Inference and dating of cacti association events in the Saccharomycotina

The number of independent events of cacti association were inferred by performing an ancestral state reconstruction using a continuous-time Markov model for discrete trait evolution implemented in Mr Bayes ^130^. A simplified workflow implemented in the R environment was followed ^131,132^. Only species classified as strictly cactophilic (Table S1) were considered as exhibiting the trait (cactophily). Transient species were considered as not having the trait. The estimated times for the emergence of cactophily were inferred according to a relaxed molecular clock analyses of the subphylum Saccharomycotina ^51^.

### Machine learning

To assess whether cactophilic species can be classified based on physiological and/or genomic traits, we used physiological data (for 893/1,154 strains) and genomic data (functional KEGG annotations) for 1,154 yeast strains ^51^. Both strictly and transiently cactophilic species were used and classified as “1” (meaning having the trait). All the remaining species in the dataset were classified as “0” (meaning lacking the trait). We trained a machine learning algorithm built by an XGBoost (1.7.3) random forest classifier (XGBRFClassifier()) with the parameters “ max_depth=12, n_estimators=100, use_label_encoder =False, eval_metric=’mlogloss’, n_jobs = 8” on 90% of the data, and we used the remaining 10% for cross-validation, using RepeatedStratifiedKFold from sklearn.model_selection (1.2.1) ^133,134^. We used RepeatedStratifiedKFold to generate accuracy measures. We used the cross_val_predict() function from Sci-Kit Learn to generate the confusion matrixes; these matrices show the numbers of strains correctly predicted to be cactophilic or non cactophilic (True Positives and True Negatives, respectively) and incorrectly predicted (False Positives, which are predicted to be cactophilic but are not; and False Negatives, which are not predicted to be, but are cactophilic). Top features were automatically generated by the XGBRFClassifier using Gini importance, which uses node impurity (the amount of variance in growth on a given carbon source for strains that either have or do not have this trait/feature). This process was repeated for 20 runs with 52 (or 31 for the analysis excluding transiently cactophilic species) randomly selected non-cactophilic species for each run, and then the averages of each result were used in the final confusion matrixes and feature importance graphs.

### Gene family evolution analyses

To find genes that are specific to or have expanded in cactophilic clades, orthogroup assignment was performed with Orthofinder v.2.3.8 ^135^ using an inflation parameter of 1.5 and DIAMOND v2.0.13.151 ^136^ as the sequence aligner. Due to the high phylogenetic distance between the major strictly cactophilic clades, in order to optimize the number of orthogroups correctly assigned, this analysis was performed separately for each cactophilic group (LDT group, Pichiales and Phaffomycetales) (Table S3). Closely related non-cactophilic species were included according to the previously reported phylogeny in Opulente & LaBella, *et. al.*, 2023 ^51^. Species in which more than 30% of the genes were multi-copy were discarded. Gene family evolutionary history was inferred using GeneRax v1.1.0 ^71^, which incorporates a Maximum-likelihood and species-tree-aware method. For that, orthogroups containing more than ten sequences were aligned with MAFFT v7.402 using an iterative refinement method (L-INS-i) ^137^. Pruned species trees for each dataset were obtained from the main Saccharomycotina tree ^51^ using PHYKIT v 1.11.12 ^138^. The species trees, and alignments were subsequently used as inputs in GeneRax. Briefly, the UndatedDTL probabilistic model was used to compute the reconciliation likelihood that accounts for duplications, transfers, and losses. For simplification, the same model of sequence evolution for all gene families (LG+I+G4) during gene tree inference by GeneRax. Reconciled trees were visualized with Notung v2.9 ^139^.

Phylogenetic trees were constructed for candidate genes/gene families putatively relevant for niche adaptation.

### Evolutionary rates

To determine which genes might exhibit altered evolutionary rates in cactophilic clades/species, we used both branch-site tests of positive selection using codeml implemented in PAML ^140^ and convergent evolutionary rates analyses implemented in RER converge ^84^ . For PAML, only strictly cactophilic species were considered and only clades containing three or more cactophilic species were included. Remaining cactophilic (strictly or transient) species were excluded. In this way, five datasets (see Fig. S3) were considered including members of different lineages (two subclades within the Phaffomycetales: *Starmera* and *Phaffomyces*; two within the Pichiales: *Pichia* A and *Pichia* B; and one within Trigonopsidales: *Tortispora*). In *Tortispora* we also included *T. agaves* because, despite not being associated with Cactaceae species, it is associated with plants with similar characteristics (*Agave* spp.) ^33^. For all these five datasets, closely related species belonging to each of the families were included based on the species phylogeny based on ^51^ (Fig. S3). Next, selection of orthogroups using Orthofinder v.2.3.8 was performed for all the species included in the five datasets. Clustering of sequences was based on protein sequence similarity and calculated using DIAMOND v2.0.13.151 using an inflation parameter of 1.5. Single copy orthologs (SCOs) present in all species in each dataset were selected. To avoid masking nonparallel signatures of positive selection, each dataset was separately analysed in PAML ^140^ using the branch-site model ^141^ and considering the branch leading to the cactophilic clade as the foreground branch. The likelihood of a gene being under positive selection was evaluated through a likelihood ratio test [LRT: 2x(ln_H1_-ln_H0_)] ^141^ using a p-value threshold of <0.001 (chi-square test with one degree of freedom; critical value = 10.8276). To avoid cases where genes under positive selection in the foreground branch were also under positive selection in closely related species (background branches), branch-site tests were performed in the same way but considering the sister clade as the foreground branch (Fig. S3) Genes for which a significant signal for positive selection was also obtained for the sister clade (p-value threshold of <0.05) were excluded. In this way, a conservative approach was employed, only considering genes for which a strong signal for positive selection was found specifically in the cactophilic branches. We further examined which genes were under positive selection in more than one clade. We assessed whether convergent amino acid substitutions occurred in genes that showed evidence of positive selection in three distinct clades (six in total: *CDA2*, *BDF1*, *PET112*, *GRS1*, *CCT3* and *SPN1*) using PCOC ^142^.

To further investigate fingerprints of convergence in evolutionary rates, we used RER converge with an extended dataset, in order to include more than one cactophilic clade/species per dataset, so that convergence could be tested (Table S3). In this case, SCOs from the Orthofinder run performed for the gene family evolution analysis were used. To increase the number of orthologs available for analysis, multi-copy orthogroups that were present in at least two species belonging to distinct cactophilic clades within each dataset were selected. Next, SCOs from each multi-copy orthogroup were pruned using OrthoSNAP v0.0.1 ^143^. Next, multiple sequence alignments were produced for each multi-copy orthogroup using MAFFT v7.402 (--localpair) and phylogenies were obtained with FastTree ^144^. We next ran OrthoSNAP with default parameters, keeping at least 50% of the species from the original dataset (50% occupancy). For each orthogroup, branch lengths were estimated on a fixed topology obtained from the Saccharomycotina species tree ^51^ by pruning the species of interest using PHYKIT v 1.11.12 ^138^. Each orthogroup was first aligned with MAFFT v7.402 (--localpair), and the best-fitting model was assessed using IQ-TREE v2.0.6. Branch lengths were determined for each orthogroup, in the fixed tree topology, using RAxML-NG v.0.9.0 ^145^ under the best protein models inferred by IQ-TREE. All phylogenies were further analysed with RER converge to find evidence of convergent evolutionary rates in cactophilic species included in each dataset.

Briefly, we tested the hypothesis of convergent evolutionary rates in the ancestral branches leading to the cactophilic clades and/or species. Only phylogenies including a minimum of two foreground species and ten species in total were considered. Genes for which a correlation ratio (Rho-correlation between relative evolutionary rate and trait) higher than 0.25 and a *p-*value (association between relative evolutionary rate and trait) lower than 0.05 were obtained were further considered as good candidates for being under convergent accelerated evolution. For those, original trees were manually checked. For Phaffomycetales, we exceptionally considered rho > 0.15 because we failed to find genes with rho > 0.25.

A detailed scheme of the entire workflow can be found in Fig. S6.

### Enrichment analyses of genes under positive selection

Gene ontology (GO) enrichment analyses were performed for the genes under positive selection in each clade (N_Starmera_=105, N_Phaffomyces_=90, N_PichiaA_= 328, N_PichiaB_= 259, N_Tortispora_=293) and for the genes under positive selection in two or more clades (N=140).

First, associated GO terms were obtained for all genes using eggNOG-mapper ^146^. For in-clade analyses, enrichment analyses were performed using *P. cactophila* (for Pichia A and PichiaB datasets), *Starmera amethionina* (for *Starmera* dataset), *Phaffomyces opuntiae* (for *Phaffomyces* dataset), and *T. caseinolytica* (for *Tortispora* dataset) whole genome annotations as the background. For the analyses involving genes under positive selection in two or more clades, *P. cactophila* genome annotations were used as the background. GO enrichment analyses were performed using the R package topGO 2.28.0 ^147^. Statistical significance was assessed using Fisher’s exact test using the “classic”, “elim”, and default “weight01” methods. Correction for multiple testing was performed the “BH” correction method. The results can be assessed in Table S7.

### Codon Usage Bias

To examine codon optimization in particular genes of cactophilic species, we calculated the gene-wise relative synonymous codon usage (gw-RSCU), implemented in Biokit v0.0.9 ^105^. This metric was shown to correlate with the tRNA adaptation index (tAI) ^105^, which measures the translation efficiency by considering both codon optimization and the intracellular concentration of tRNA molecules ^148^. The gw-RSCU was calculated by determining the mean relative synonymous codon usage value for all codons in each gene in the genome based on their genome-wide RSCU values. We ranked the genes with the highest gw-RSCU values (subtracting the standard deviation to the gw-RSCU mean value) and looked at the genes falling into the 95^th^ percentile and above (Table S8). Next, gene functions that were relevant for the cactophilic lifestyle were selected, and their gw-RSCU values were inspected in non-cactophilic closest related species. Briefly, a local BLASTp was used to find the putative orthologs by considering a protein identity of > 40%. The top hit was considered to correspond to the orthologous gene; however, whenever multiple hits with similar protein sequence identity were found, the one with the highest rank was considered. The percentile ranking for *CDA2* was determined using the R package *dplyr* ^149^.

## Acknowledgements

We thank members of the Rokas Lab, Hittinger Lab, and members of the Y1000+ project (http://y1000plus.org) for helpful discussions. This work was performed using resources contained within the Advanced Computing Center for research and Education at Vanderbilt University in Nashville, TN.

## Funding

National Science Foundation Grant DEB-2110403 (CTH)

National Science Foundation Grant DEB-2110404 (AR)

DOE Great Lakes Bioenergy Research Center, funded by BER Office of Science Grant DE-SC0018409 (CTH)

USDA National Institute of Food and Agriculture Hatch Project 1020204 (CTH)

H. I. Romnes Faculty Fellow, supported by the Office of the Vice Chancellor for Research and Graduate Education with funding from the Wisconsin Alumni Research Foundation (CTH)

National Institutes of Health/National Institute of Allergy and Infectious Diseases Grant R01 AI153356 (AR)

Burroughs Wellcome Fund (AR)

Research supported by the National Key R&D Program of China Grant 2022YFD1401600 (XXS)

National Science Foundation for Distinguished Young Scholars of Zhejiang Province Grant LR23C140001 (XXS)

Fundamental Research Funds for the Central Universities Grant 226-2023-00021 (XXS) National Institutes of Health Grant T32 HG002760-16 (JFW)

National Science Foundation Grant Postdoctoral Research Fellowship in Biology 1907278 (JFW)

JLS is a Howard Hughes Medical Institute Awardee of the Life Sciences Research Foundation.

