## Supplementary Figures and Supplementary Table Legends for "Diverse signatures of convergent evolution in cacti-associated yeasts"

**Supplemental material:**

**Fig. S1. Multiple independent evolution of cactophily across the subphylum Saccharomycotina.** Predicted number of independent emergences of cacti association according to ancestral state reconstruction by Mr Bayes.

**Fig. S2. Cactophilic plant cell wall-degrading proteins that underwent duplication or HGT events localize at the extracellular space.** Deeploc analyses showing predicted localization of proteins involved in plant cell wall degradation for which either duplication or HGT events were detected.

**Fig. S3**. **PAML datasets showing the species trees used and the control tests in sister clades.** Branches selected for the analyses are shown (ω) in different colors (green for the foreground/cactophilic lineages and red for the background/non-cactophilic lineages). Only the genes for which no evidence of positive selection on the background lineages were considered for further analyses (please see Materials and Methods section).

**Fig. S4.** ***CDA2*** **gw-RSCU percentile rank in cactophilic species**. Cactophilic species belonging to *Phaffomyces*, *Starmera*, *Pichia* and *Myxozyma* genera and their respective non-cactophilic closest relatives (identified as "out") were inspected.

**Fig. S5**. **Ecological associations of cactophilic and their closest relatives**. Species that have been isolated from clinical contexts, and are emerging opportunistic pathogens, are highlighted. The ecological information presented was obtained from the CBS database and available literature according to the substrate of isolation of the type strain.

**Fig. S6. Workflow of RER converge analyses.**

**Table S1. List of cactophilic species understudy.** Selection of cacti-associated yeasts according to the literature (relevant references are shown). Classification of cactophilic yeasts (strict or transient), geography and substrate of origin is shown for the type strain.

**Table S2. Raw machine learning results.** Proportion of presence in cactophilic and non-cactophilic species are shown for the top 100 most important features.

**Table S3. List of species used for gene family analyses and RER converge analyses.**

**Table S4. Gene family evolution results.** Duplication events in at least one species belonging to yeast cactophilic group inspected were considered. The respective functional annotations obtained from EggNOG are shown. The table below shows the functional annotations in common between the three groups understudy.

**Table S5. RER converge results.** Genes that underwent accelerated and decelerated evolution as well as their putative functions (according to SGD).

**Table S6**. **Results of positive selection analyses from branch-site tests using PAML for the five cactophilic datasets understudy (*Tortispora*, *Pichia* A, *Pichia* B, *Starmera* and *Phaffomyces*).** Putative genes from each orthogroup were identified after a BLASTp search against the nr NCBI database using Saccharomyces (taxid:4930) as the reference. When no hit was obtained for *S. cerevisiae*, a BLASTp against the entire nr database was performed instead.

**Table S7. GO term enrichment results.** GO terms were obtained for all genes using eggNOG-mapper for genes under positive selection in each clade and for genes under positive selection in two or more clades.

**Table S8. Top-ranked (95$^{th}$ percentile) genes for gw-RSCU in cactophilic species.**

**Fig. S1**

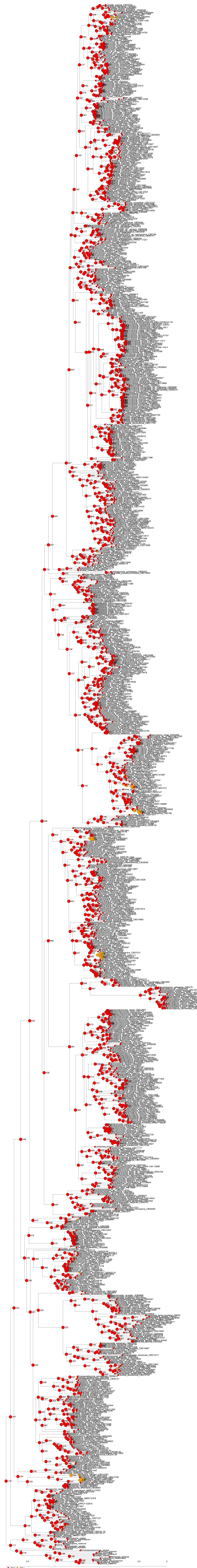

**Fig. S1**

### Fig. S2

Download prediction results: CSV Summary (/services/DeepLoc-2.0/tmp/64A9573700004F884151921D/results_64A9573700004F884151921D.csv)

Probability thresholds

#### Summary of 8 predicted sequences

Table of predicted subcelullar localizations. Use the Instructions page for more detailed description of the output page.

**Phaffomyces_opuntiae_g001049.m1**
**Predicted localizations:** Extracellular
**Predicted signals:** Signal peptide

| Localization | Cytoplasm | Nucleus | Extracellular | Cell membrane | Mitochondrion | Plastid | Endoplasmic reticulum | Lysosome/Vacuole | Golgi apparatus | Peroxisome |
|---|---|---|---|---|---|---|---|---|---|---|
| **Probability** | 0.2096 | 0.0618 | 0.8641 | 0.0642 | 0.0866 | 0.0203 | 0.5346 | 0.3550 | 0.2089 | 0.0029 |

**Sorting Signal Importance. Donwload:** PNG (/services/DeepLoc-2.0/tmp/64A9573700004F884151921D/alpha_phaffomyces_opuntiae_g001049m1.png) **/** CSV (/services/DeepLoc-2.0/tmp/64A9573700004F884151921D/alpha_phaffomyces_opuntiae_g001049m1.csv)

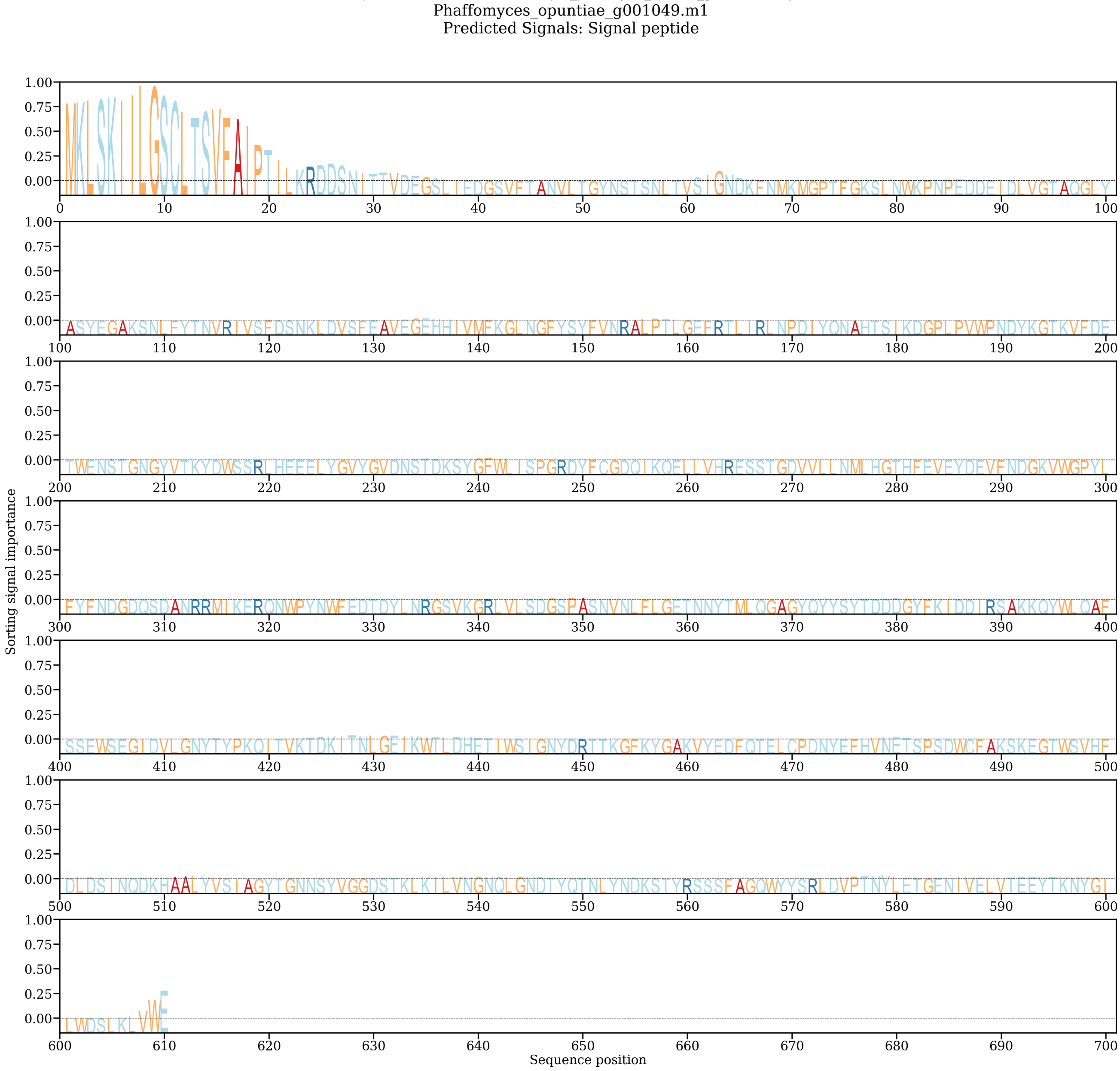


**Phaffomyces_opuntiae_g002229.m1**
**Predicted localizations:** Extracellular
**Predicted signals:** Signal peptide

| Localization | Cytoplasm | Nucleus | Extracellular | Cell membrane | Mitochondrion | Plastid | Endoplasmic reticulum | Lysosome/Vacuole | Golgi apparatus | Peroxisome |
|---|---|---|---|---|---|---|---|---|---|---|
| **Probability** | 0.4458 | 0.1420 | 0.8031 | 0.4817 | 0.1344 | 0.0394 | 0.4889 | 0.3267 | 0.1224 | 0.0085 |

**Sorting Signal Importance. Donwload:** PNG (/services/DeepLoc-2.0/tmp/64A9573700004F884151921D/alpha_phaffomyces_opuntiae_g002229m1.png) **/** CSV (/services/DeepLoc-2.0/tmp/64A9573700004F884151921D/alpha_phaffomyces_opuntiae_g002229m1.csv)

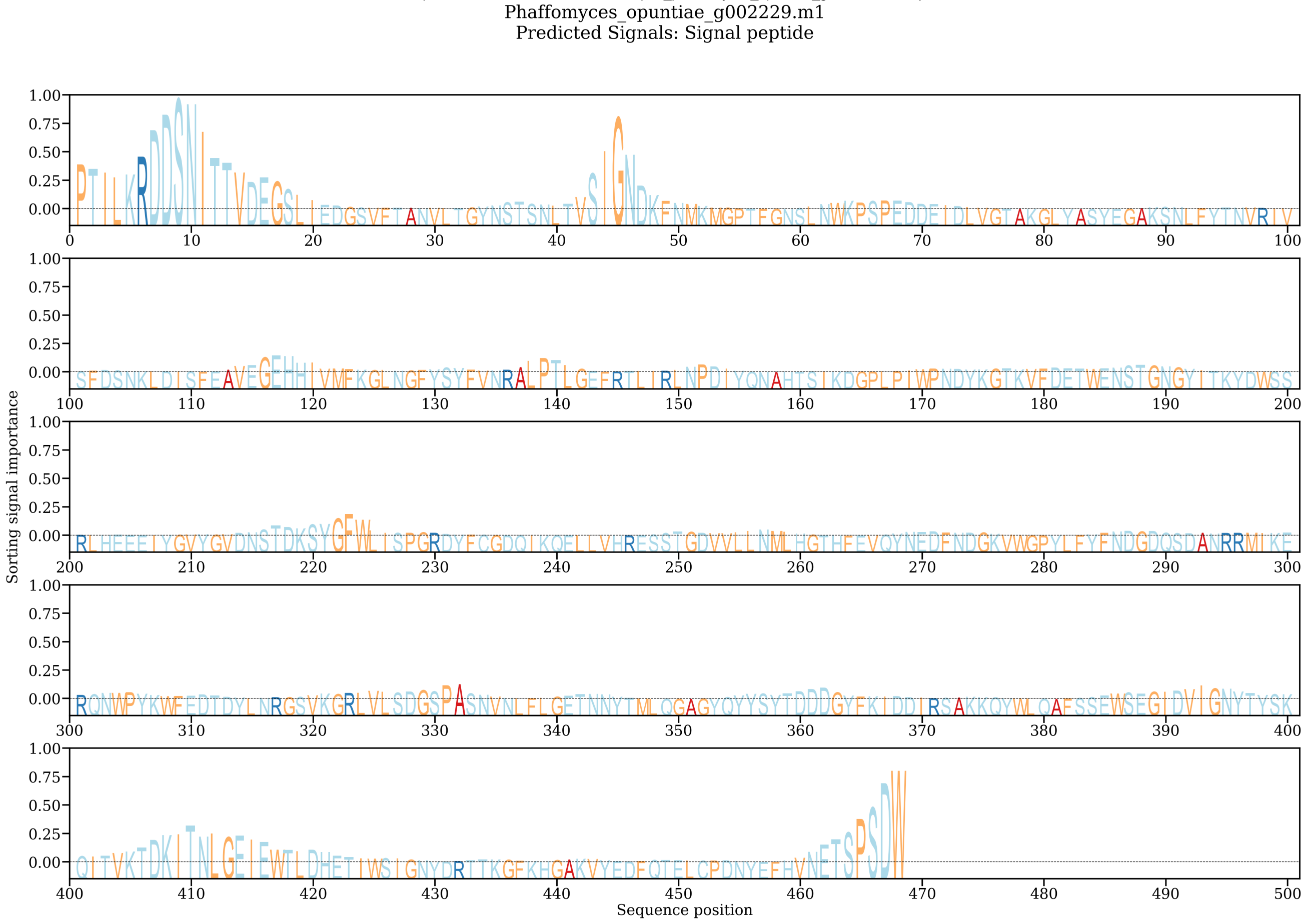


**Candida_coquimbonensis_g003482.m1**

**Predicted localizations:** Extracellular, Endoplasmic reticulum

**Predicted signals:** Signal peptide

| Localization | Cytoplasm | Nucleus | Extracellular | Cell membrane | Mitochondrion | Plastid | Endoplasmic reticulum | Lysosome/Vacuole | Golgi apparatus | Peroxisome |
|---|---|---|---|---|---|---|---|---|---|---|
| **Probability** | 0.2701 | 0.1194 | 0.8169 | 0.1238 | 0.0529 | 0.0152 | 0.6108 | 0.4708 | 0.2705 | 0.0015 |

**Sorting Signal Importance. Donwload:** PNG (/services/DeepLoc-2.0/tmp/64A9573700004F884151921D/alpha_candida_coquimbonensis_g003482m1.png) **/** CSV (/services/DeepLoc-2.0/tmp/64A9573700004F884151921D/alpha_candida_coquimbonensis_g003482m1.csv)

Candida_coquimbonensis_g003482.m1
Predicted Signals: Signal peptide

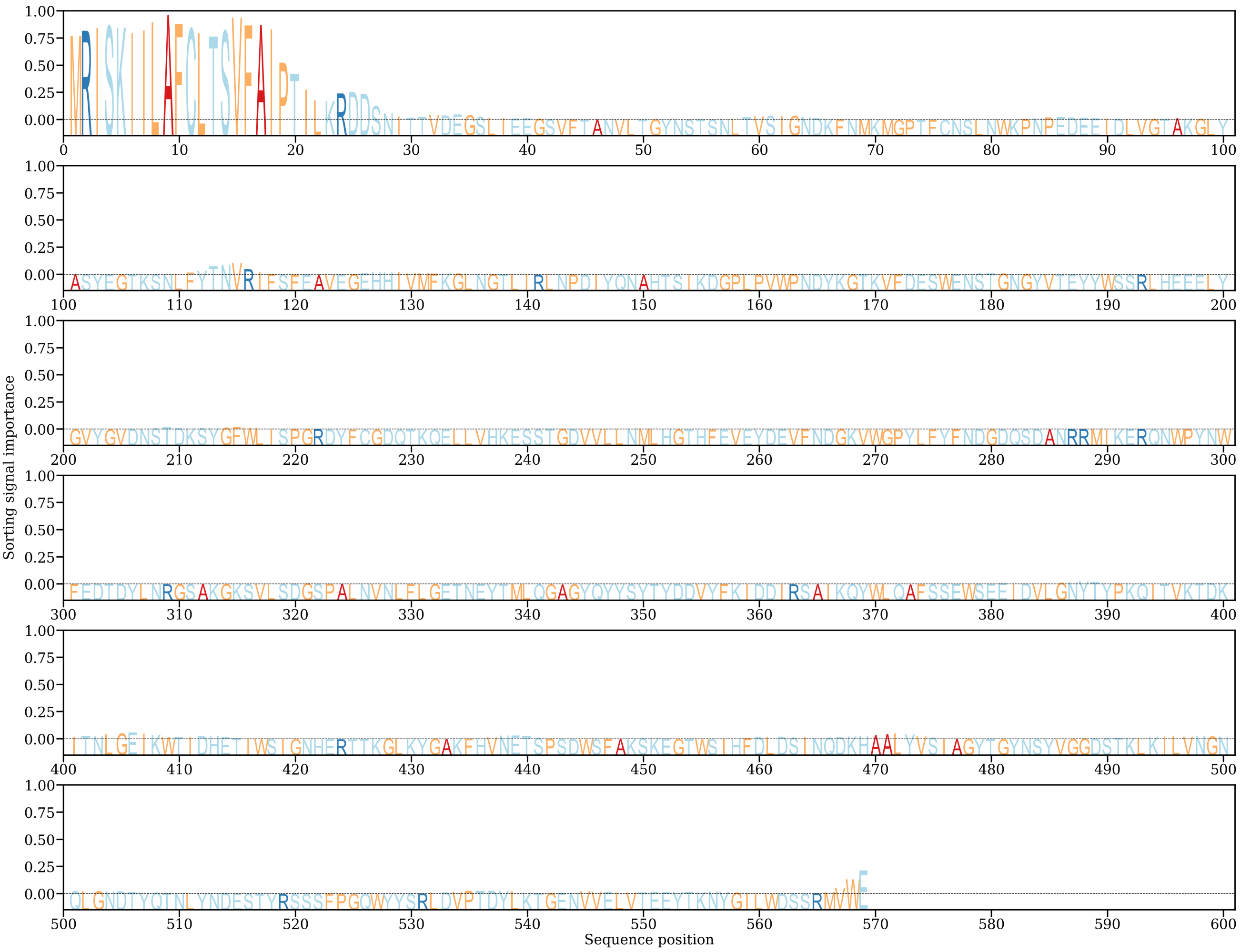


**Candida_coquimbonensis_g004454.m1**
**Predicted localizations:** Extracellular
**Predicted signals:** Signal peptide

| Localization | Cytoplasm | Nucleus | Extracellular | Cell membrane | Mitochondrion | Plastid | Endoplasmic reticulum | Lysosome/Vacuole | Golgi apparatus | Peroxisome |
|---|---|---|---|---|---|---|---|---|---|---|
| **Probability** | 0.1693 | 0.0654 | 0.8567 | 0.0685 | 0.0682 | 0.0169 | 0.5151 | 0.3625 | 0.2297 | 0.0033 |

**Sorting Signal Importance. Donwload:** PNG (/services/DeepLoc-2.0/tmp/64A9573700004F884151921D/alpha_candida_coquimbonensis_g004454m1.png) **/** CSV (/services/DeepLoc-2.0/tmp/64A9573700004F884151921D/alpha_candida_coquimbonensis_g004454m1.csv)

Candida_coquimbonensis_g004454.m1
Predicted Signals: Signal peptide

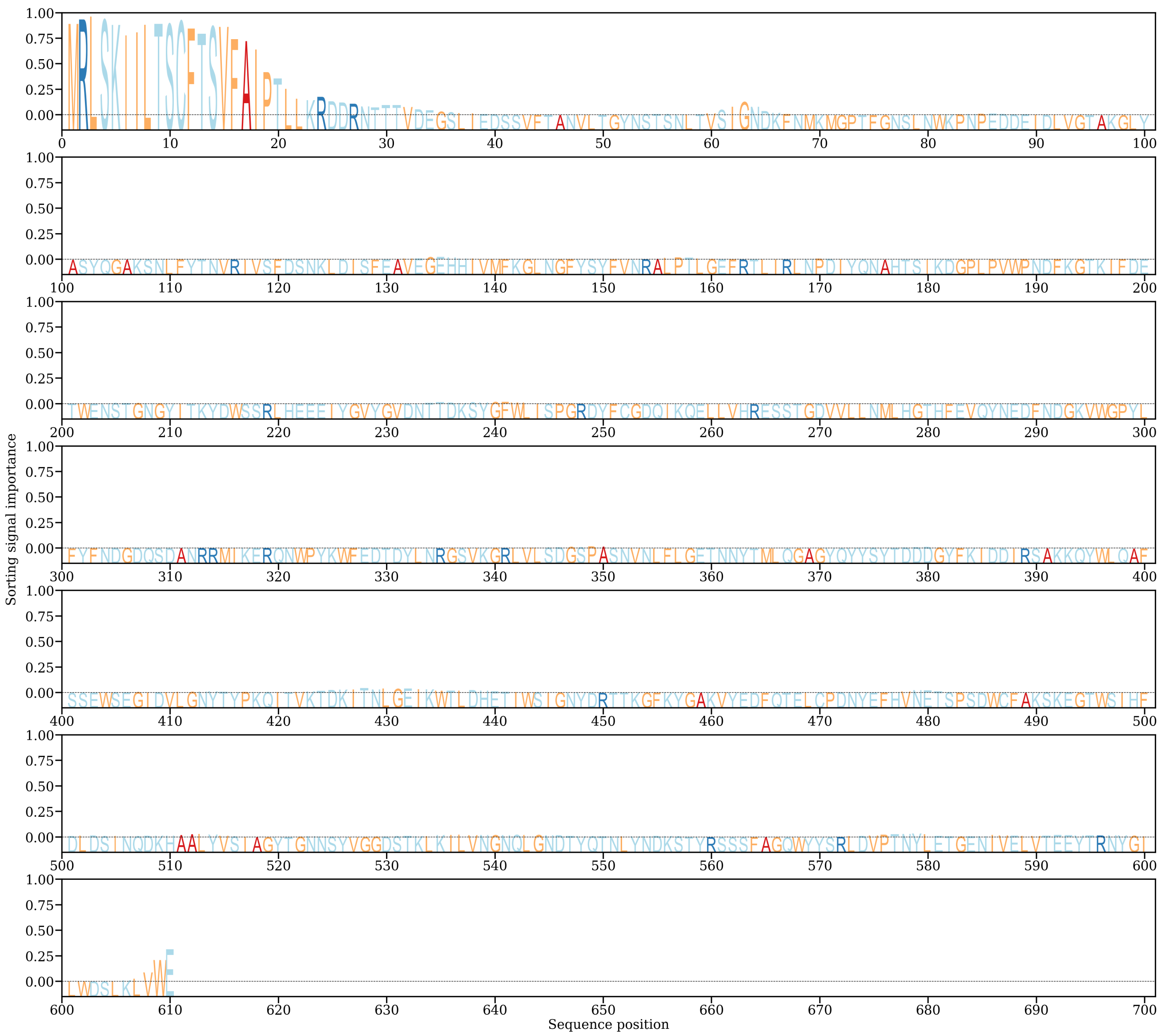

**Phaffomyces_antillensis_g005279.m1**
**Predicted localizations:** Extracellular
**Predicted signals:** Signal peptide

| Localization | Cytoplasm | Nucleus | Extracellular | Cell membrane | Mitochondrion | Plastid | Endoplasmic reticulum | Lysosome/Vacuole | Golgi apparatus | Peroxisome |
|---|---|---|---|---|---|---|---|---|---|---|
| **Probability** | 0.1708 | 0.0644 | 0.8661 | 0.0477 | 0.0701 | 0.0148 | 0.4198 | 0.2902 | 0.2075 | 0.0023 |

**Sorting Signal Importance. Donwload:** PNG (/services/DeepLoc-2.0/tmp/64A9573700004F884151921D/alpha_phaffomyces_antillensis_g005279m1.png) **/** CSV (/services/DeepLoc-2.0/tmp/64A9573700004F884151921D/alpha_phaffomyces_antillensis_g005279m1.csv)

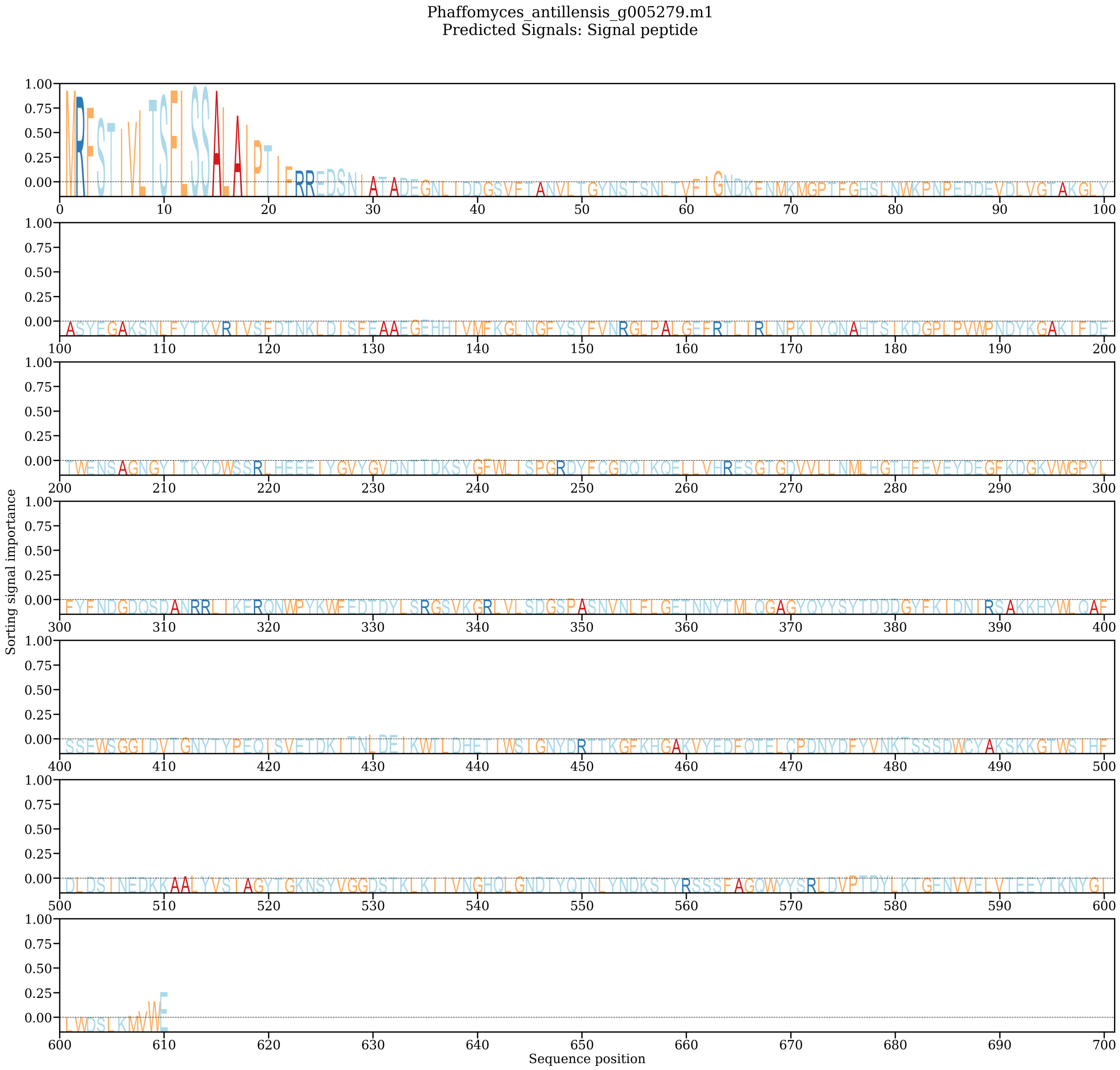


**Phaffomyces_antillensis_g003707.m1**
**Predicted localizations:** Extracellular
**Predicted signals:** Signal peptide

| Localization | Cytoplasm | Nucleus | Extracellular | Cell membrane | Mitochondrion | Plastid | Endoplasmic reticulum | Lysosome/Vacuole | Golgi apparatus | Peroxisome |
|---|---|---|---|---|---|---|---|---|---|---|
| **Probability** | 0.1838 | 0.0752 | 0.8814 | 0.0542 | 0.0537 | 0.0082 | 0.4310 | 0.2891 | 0.2099 | 0.0016 |

**Sorting Signal Importance. Donwload:** PNG (/services/DeepLoc-2.0/tmp/64A9573700004F884151921D/alpha_phaffomyces_antillensis_g003707m1.png) **/** CSV (/services/DeepLoc-2.0/tmp/64A9573700004F884151921D/alpha_phaffomyces_antillensis_g003707m1.csv)

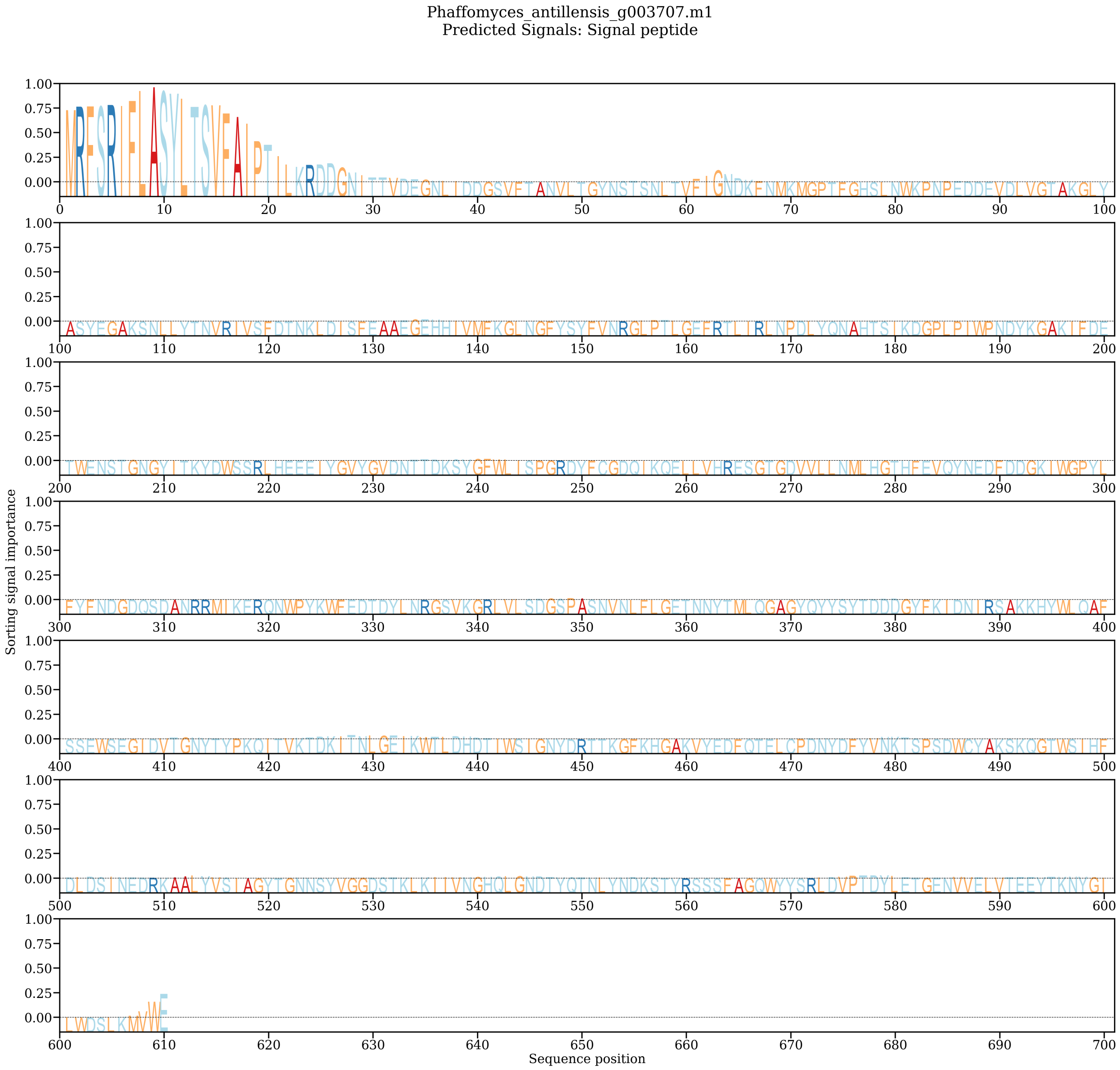


**Pectin_lyase_Pichia_eremophila**
**Predicted localizations:** Extracellular
**Predicted signals:** Signal peptide

| Localization | Cytoplasm | Nucleus | Extracellular | Cell membrane | Mitochondrion | Plastid | Endoplasmic reticulum | Lysosome/Vacuole | Golgi apparatus | Peroxisome |
|---|---|---|---|---|---|---|---|---|---|---|
| **Probability** | 0.2170 | 0.2693 | 0.6924 | 0.1076 | 0.2783 | 0.0520 | 0.2464 | 0.2624 | 0.2556 | 0.0007 |

**Sorting Signal Importance. Donwload:** PNG (/services/DeepLoc-2.0/tmp/64A9573700004F884151921D/alpha_pectin_lyase_pichia_eremophila.png) **/** CSV (/services/DeepLoc-2.0/tmp/64A9573700004F884151921D/alpha_pectin_lyase_pichia_eremophila.csv)

Pectin_lyase_Pichia_eremophila
Predicted Signals: Signal peptide

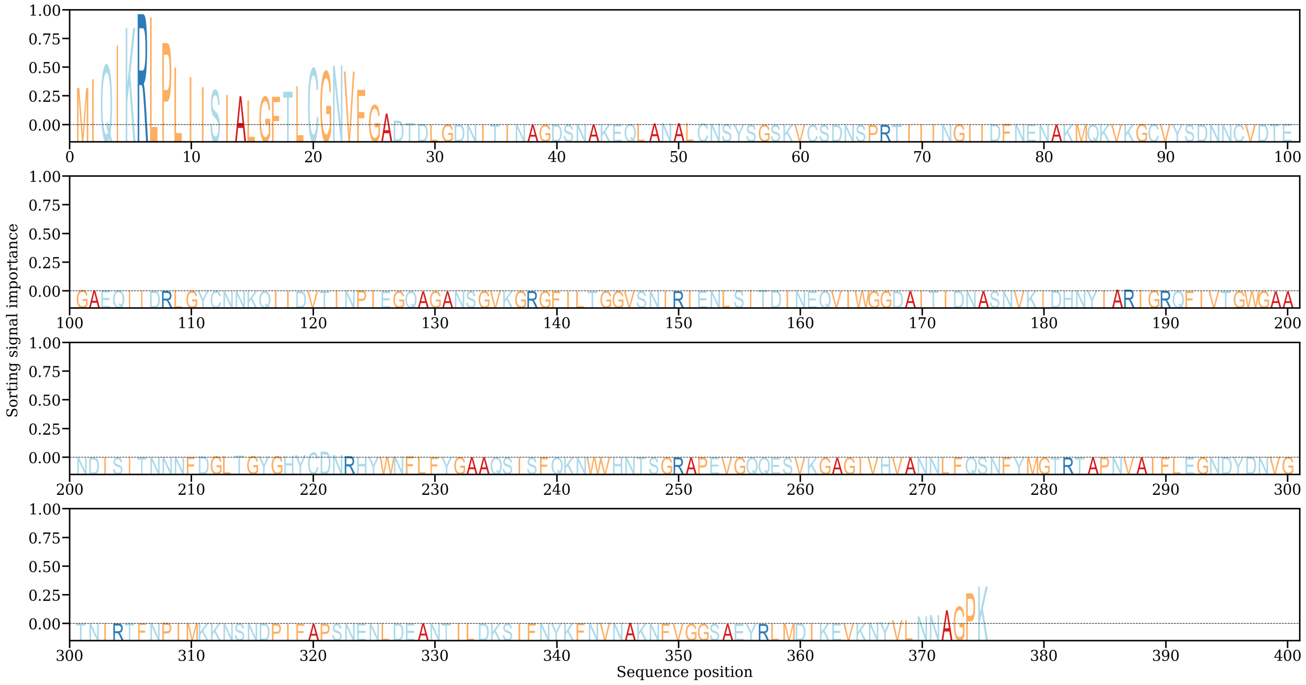


**Pectin_lyase_Pichia_kluyveri**
**Predicted localizations:** Extracellular
**Predicted signals:** Signal peptide

| Localization | Cytoplasm | Nucleus | Extracellular | Cell membrane | Mitochondrion | Plastid | Endoplasmic reticulum | Lysosome/Vacuole | Golgi apparatus | Peroxisome |
|---|---|---|---|---|---|---|---|---|---|---|
| **Probability** | 0.2222 | 0.3077 | 0.7363 | 0.0696 | 0.2125 | 0.0336 | 0.2847 | 0.3683 | 0.3591 | 0.0019 |

**Sorting Signal Importance. Donwload:** PNG (/services/DeepLoc-2.0/tmp/64A9573700004F884151921D/alpha_pectin_lyase_pichia_kluyveri.png) **/** CSV (/services/DeepLoc-2.0/tmp/64A9573700004F884151921D/alpha_pectin_lyase_pichia_kluyveri.csv)

Pectin_lyase_Pichia_kluyveri
Predicted Signals: Signal peptide

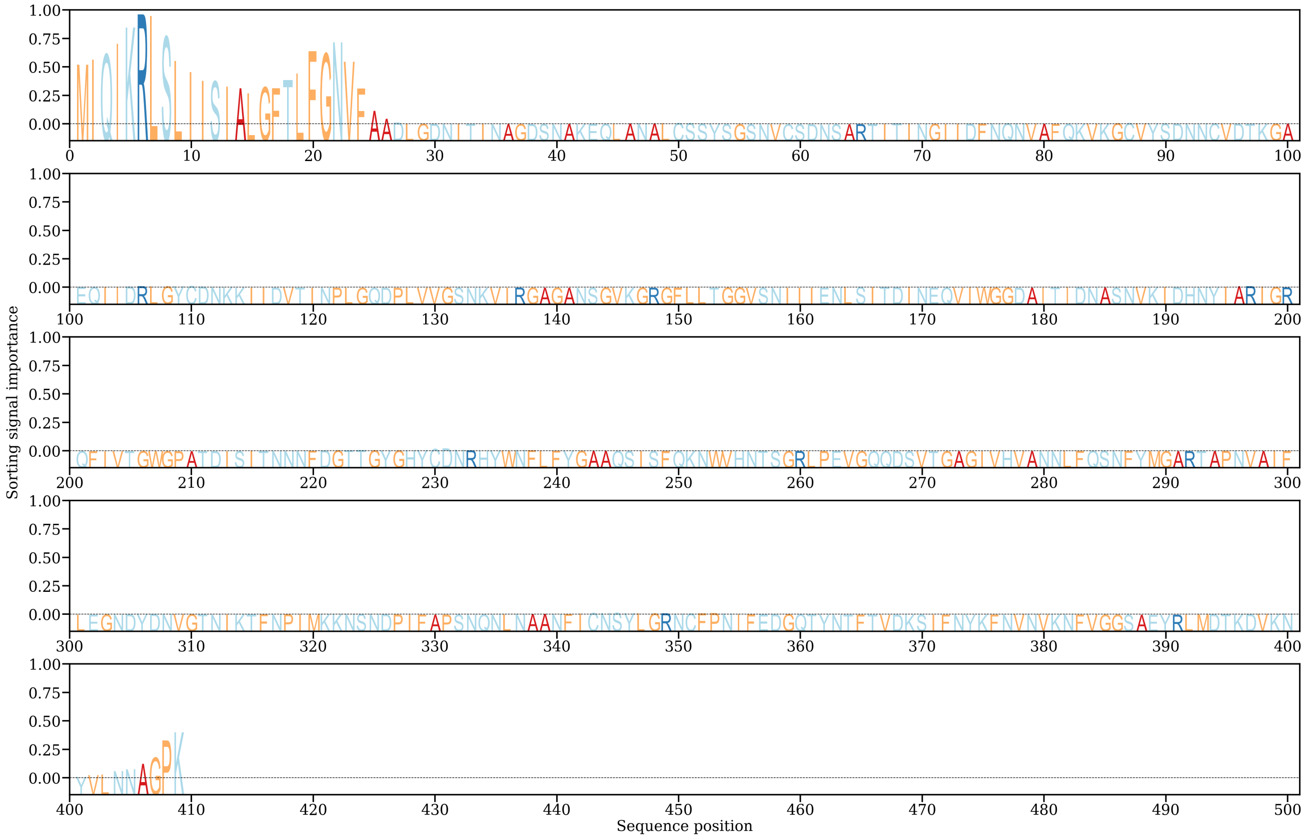


Found device of type cuda: Tesla V100-PCIE-16GB Loading model... Running model on device of type cuda: Tesla V100-PCIE-16GB Model loaded in 52.92s Attention plotted in 63.33s Model run time: 1.44s Finished prediction in 121.18s

Download prediction results: CSV Summary (/services/DeepLoc-2.0/tmp/64A53F1F000077FFC33E38A4/results_64A53F1F000077FFC33E38A4.csv)

Probability thresholds

#### Summary of 4 predicted sequences

Table of predicted subcelullar localizations. Use the Instructions page for more detailed description of the output page.

**Dipodascus_australiensis_g006729.m1**
**Predicted localizations:** Extracellular
**Predicted signals:** Signal peptide

| Localization | Cytoplasm | Nucleus | Extracellular | Cell membrane | Mitochondrion | Plastid | Endoplasmic reticulum | Lysosome/Vacuole | Golgi apparatus | Peroxisome |
|---|---|---|---|---|---|---|---|---|---|---|
| **Probability** | 0.2265 | 0.0464 | 0.9536 | 0.0571 | 0.0867 | 0.0025 | 0.2312 | 0.2042 | 0.2017 | 0.0003 |

**Sorting Signal Importance. Donwload:** PNG (/services/DeepLoc-2.0/tmp/64A53F1F000077FFC33E38A4/alpha_dipodascus_australiensis_g006729m1.png) **/** CSV (/services/DeepLoc-2.0/tmp/64A53F1F000077FFC33E38A4/alpha_dipodascus_australiensis_g006729m1.csv)

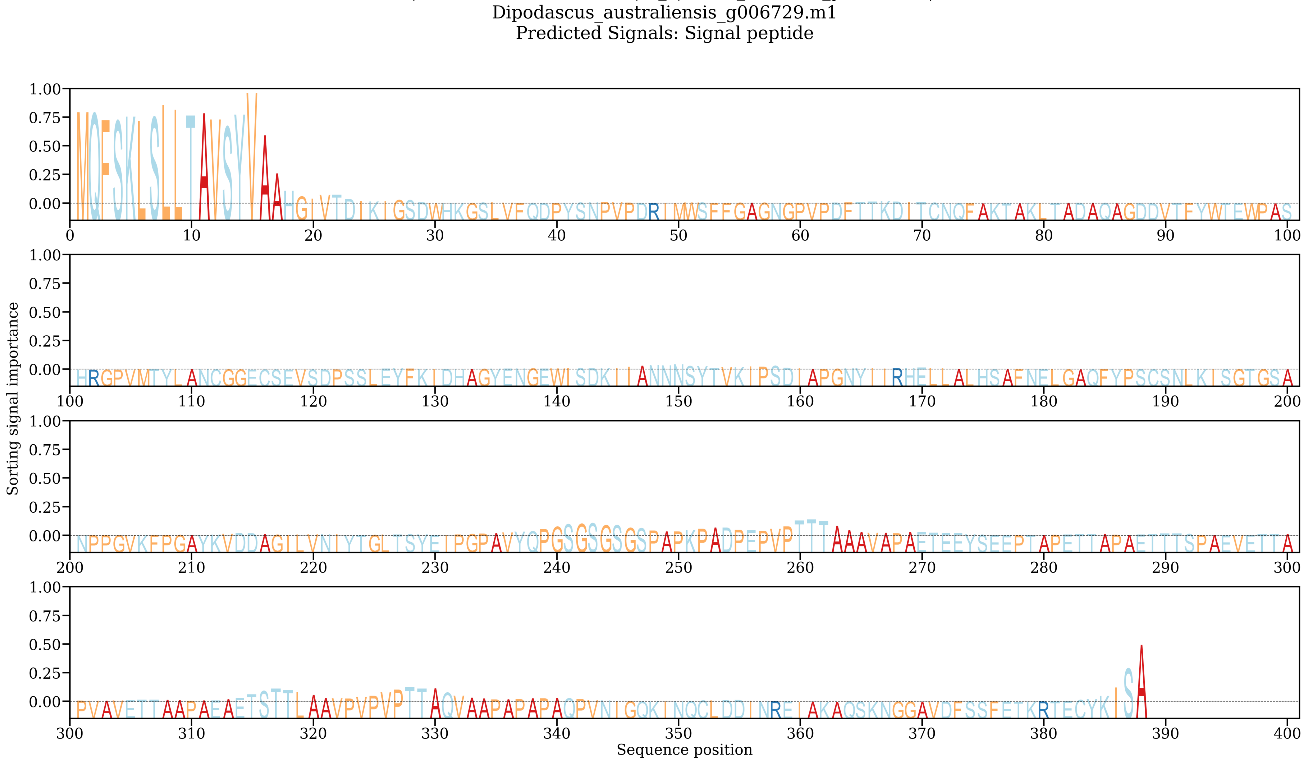


**Dipodascus_australiensis_g000485.m1**
**Predicted localizations:** Extracellular
**Predicted signals:** Signal peptide

| Localization | Cytoplasm | Nucleus | Extracellular | Cell membrane | Mitochondrion | Plastid | Endoplasmic reticulum | Lysosome/Vacuole | Golgi apparatus | Peroxisome |
|---|---|---|---|---|---|---|---|---|---|---|
| **Probability** | 0.2864 | 0.0439 | 0.9524 | 0.0646 | 0.0873 | 0.0018 | 0.2092 | 0.1809 | 0.1951 | 0.0002 |

**Sorting Signal Importance. Donwload:** PNG (/services/DeepLoc-2.0/tmp/64A53F1F000077FFC33E38A4/alpha_dipodascus_australiensis_g000485m1.png) **/** CSV (/services/DeepLoc-2.0/tmp/64A53F1F000077FFC33E38A4/alpha_dipodascus_australiensis_g000485m1.csv)

Dipodascus_australiensis_g000485.m1
Predicted Signals: Signal peptide

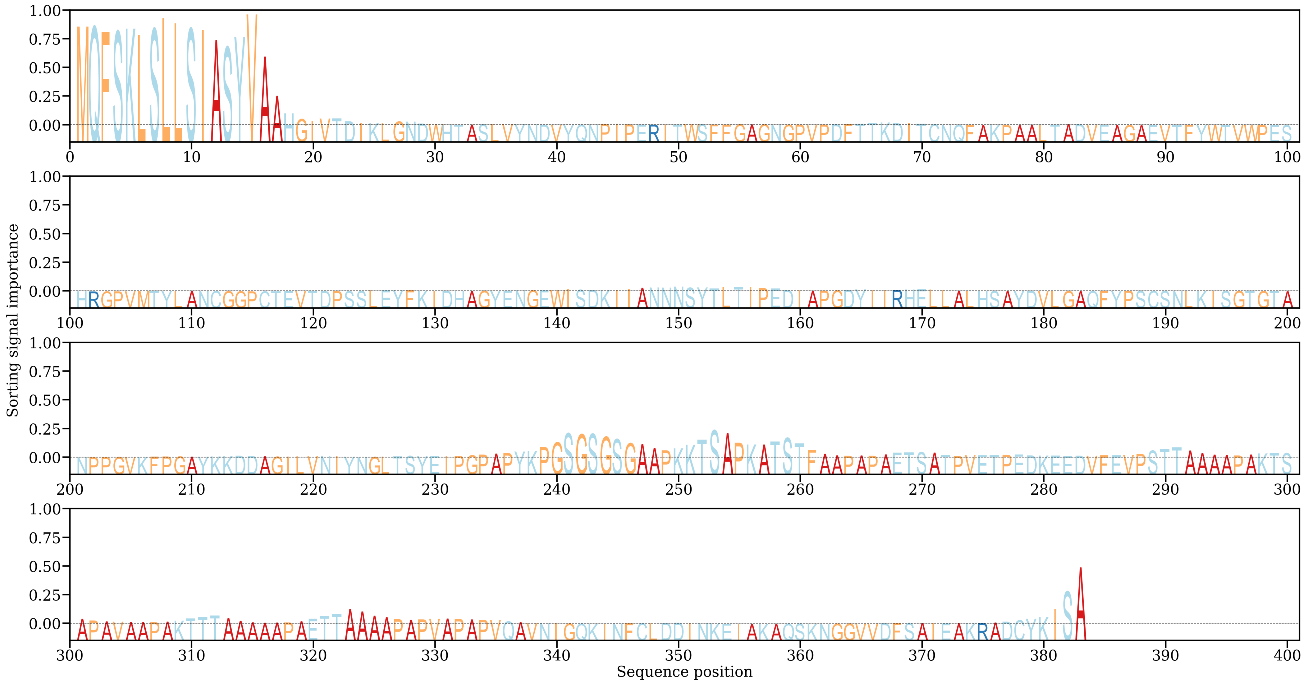


**Dipodascus_australiensis_g001311.m1**

**Predicted localizations:** Extracellular

**Predicted signals:** Signal peptide

| Localization | Cytoplasm | Nucleus | Extracellular | Cell membrane | Mitochondrion | Plastid | Endoplasmic reticulum | Lysosome/Vacuole | Golgi apparatus | Peroxisome |
|---|---|---|---|---|---|---|---|---|---|---|
| **Probability** | 0.1718 | 0.0802 | 0.9109 | 0.0583 | 0.0885 | 0.0022 | 0.3142 | 0.1608 | 0.2244 | 0.0018 |

**Sorting Signal Importance. Donwload:** PNG (/services/DeepLoc-2.0/tmp/64A53F1F000077FFC33E38A4/alpha_dipodascus_australiensis_g001311m1.png) **/** CSV (/services/DeepLoc-2.0/tmp/64A53F1F000077FFC33E38A4/alpha_dipodascus_australiensis_g001311m1.csv)

Dipodascus_australiensis_g001311.m1
Predicted Signals: Signal peptide

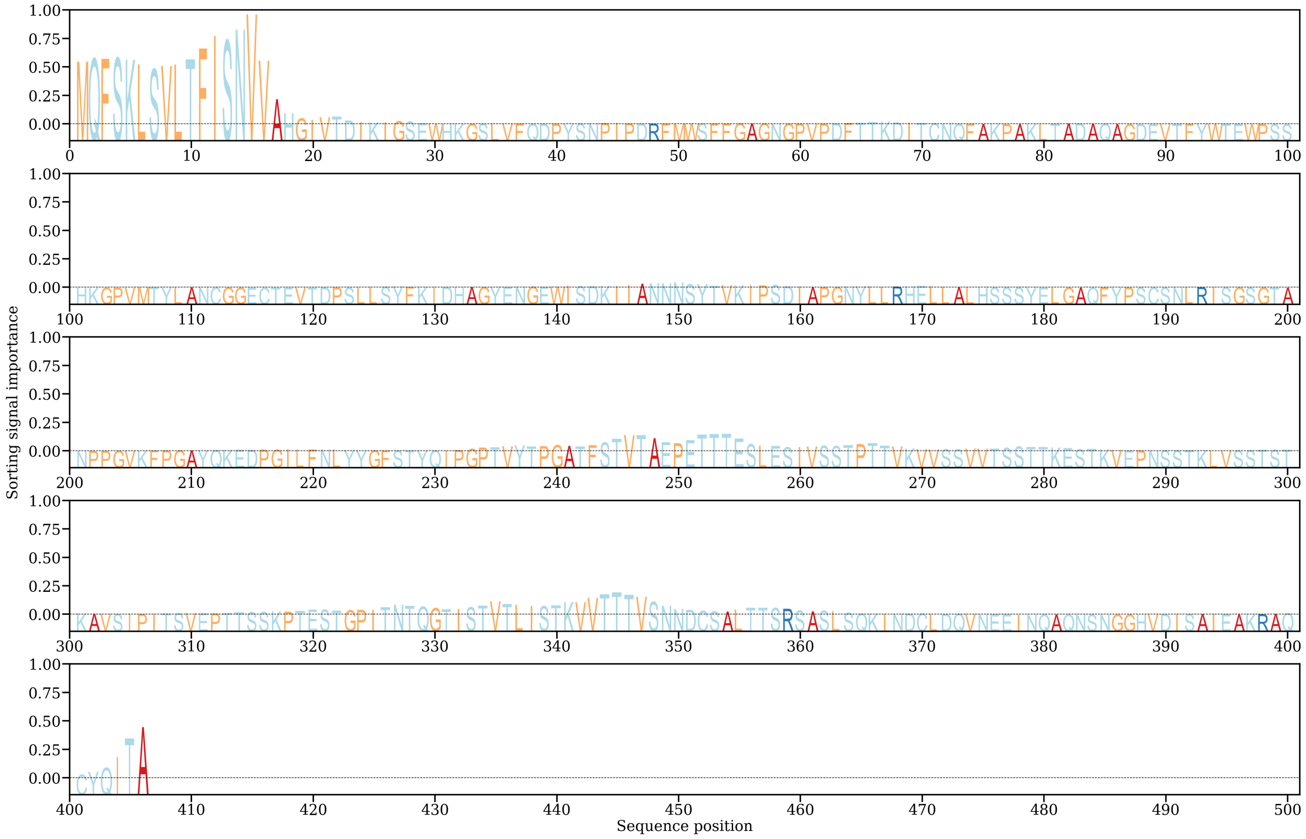

**Dipodascus_australiensis_g002066.m1**

**Predicted localizations:** Extracellular

**Predicted signals:** Signal peptide

| Localization | Cytoplasm | Nucleus | Extracellular | Cell membrane | Mitochondrion | Plastid | Endoplasmic reticulum | Lysosome/Vacuole | Golgi apparatus | Peroxisome |
|---|---|---|---|---|---|---|---|---|---|---|
| **Probability** | 0.2117 | 0.0806 | 0.9180 | 0.0646 | 0.0871 | 0.0036 | 0.3617 | 0.1814 | 0.2692 | 0.0009 |

**Sorting Signal Importance. Donwload:** PNG (/services/DeepLoc-2.0/tmp/64A53F1F000077FFC33E38A4/alpha_dipodascus_australiensis_g002066m1.png) / CSV (/services/DeepLoc-2.0/tmp/64A53F1F000077FFC33E38A4/alpha_dipodascus_australiensis_g002066m1.csv)

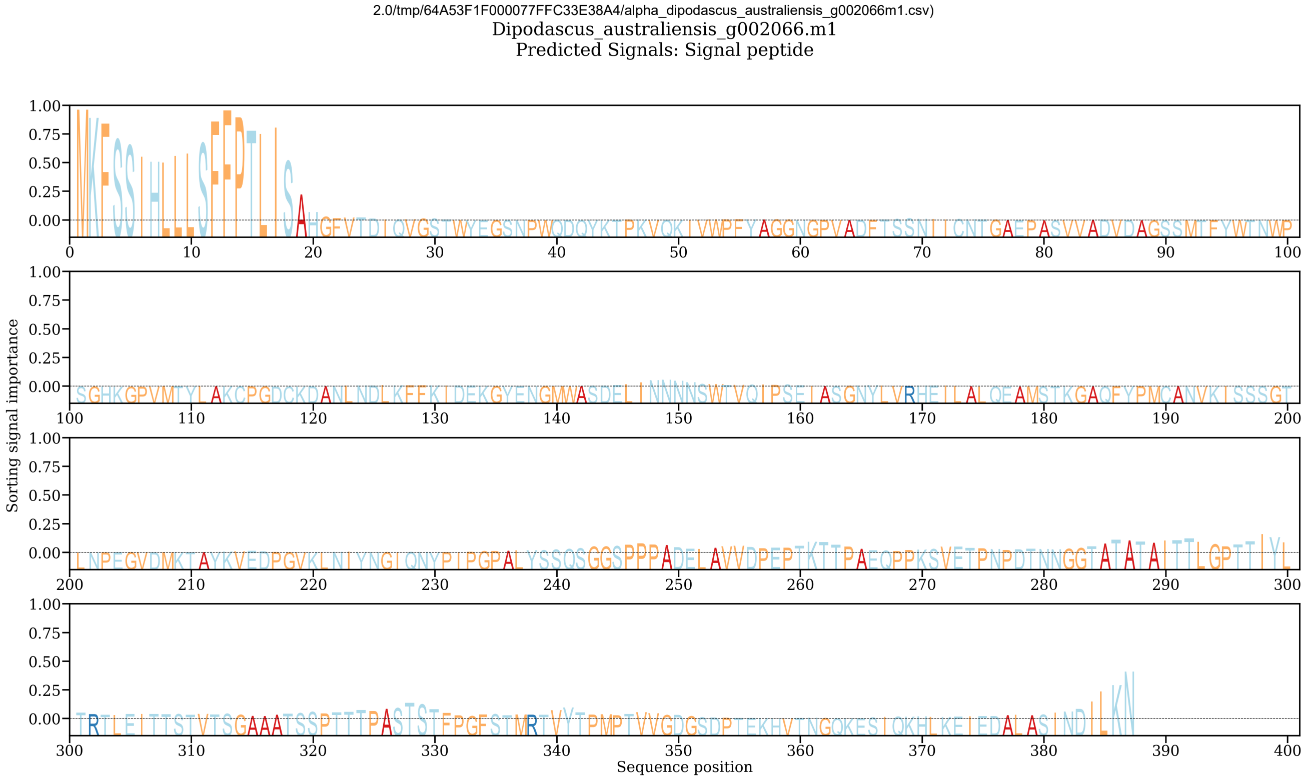


Found device of type cuda: Tesla V100-PCIE-16GB Loading model... Running model on device of type cuda: Tesla V100-PCIE-16GB Model loaded in 31.31s Attention plotted in 24.40s Model run time: 0.48s Finished prediction in 58.64s

Download prediction results: CSV Summary (/services/DeepLoc-2.0/tmp/64A6C5DD00002D2A7A92F90C/results_64A6C5DD00002D2A7A92F90C.csv)

Probability thresholds

#### Summary of 15 predicted sequences

Table of predicted subcelullar localizations. Use the Instructions page for more detailed description of the output page.

##### Starmera_pachycereana_scaf28

**Predicted localizations:** Extracellular

**Predicted signals:** Signal peptide

| Localization | Cytoplasm | Nucleus | Extracellular | Cell membrane | Mitochondrion | Plastid | Endoplasmic reticulum | Lysosome/Vacuole | Golgi apparatus | Peroxisome |
|---|---|---|---|---|---|---|---|---|---|---|
| **Probability** | 0.1911 | 0.0912 | 0.8969 | 0.1288 | 0.0533 | 0.0025 | 0.2544 | 0.2909 | 0.2916 | 0.0024 |

**Sorting Signal Importance. Donwload:** PNG (/services/DeepLoc-2.0/tmp/64A6C5DD00002D2A7A92F90C/alpha_starmera_pachycereana_scaf28.png) / CSV (/services/DeepLoc-2.0/tmp/64A6C5DD00002D2A7A92F90C/alpha_starmera_pachycereana_scaf28.csv)

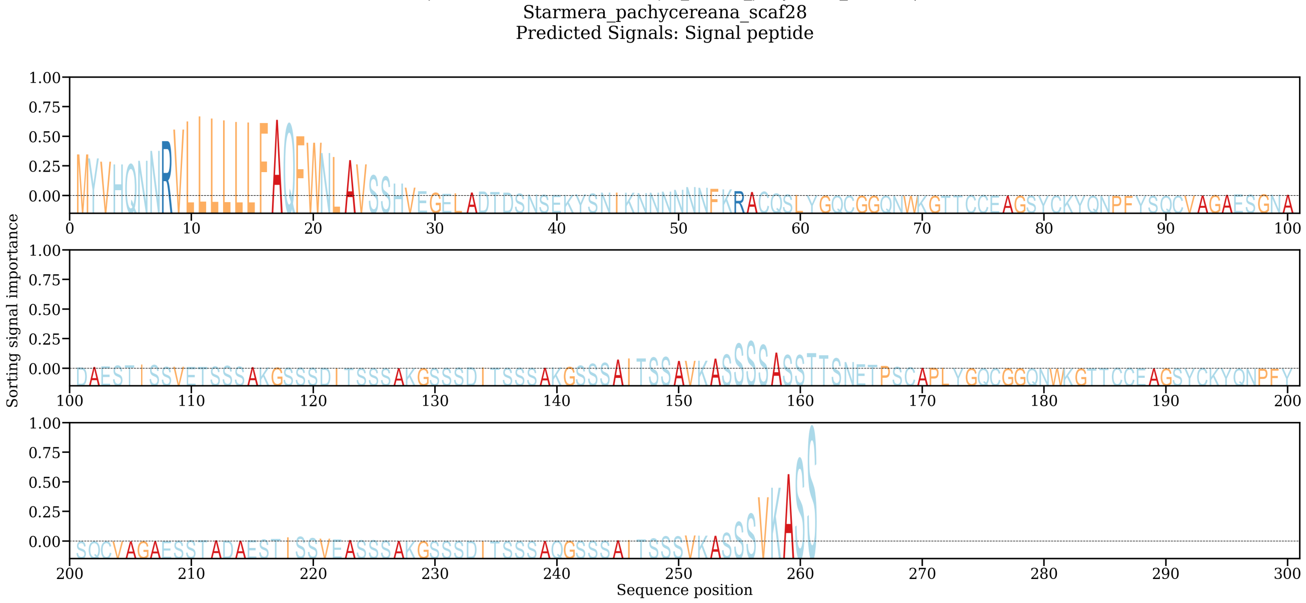


##### Starmera_pachycereana_scaf812

**Predicted localizations:** Extracellular

**Predicted signals:**

| Localization | Cytoplasm | Nucleus | Extracellular | Cell membrane | Mitochondrion | Plastid | Endoplasmic reticulum | Lysosome/Vacuole | Golgi apparatus | Peroxisome |
|---|---|---|---|---|---|---|---|---|---|---|
| **Probability** | 0.4710 | 0.1714 | 0.9060 | 0.1485 | 0.0405 | 0.0018 | 0.0291 | 0.0497 | 0.0522 | 0.0423 |

**Sorting Signal Importance. Donwload:** PNG (/services/DeepLoc-2.0/tmp/64A6C5DD00002D2A7A92F90C/alpha_starmera_pachycereana_scaf812.png) / CSV (/services/DeepLoc-2.0/tmp/64A6C5DD00002D2A7A92F90C/alpha_starmera_pachycereana_scaf812.csv)

Starmera_pachycereana_scaf812
Predicted Signals:

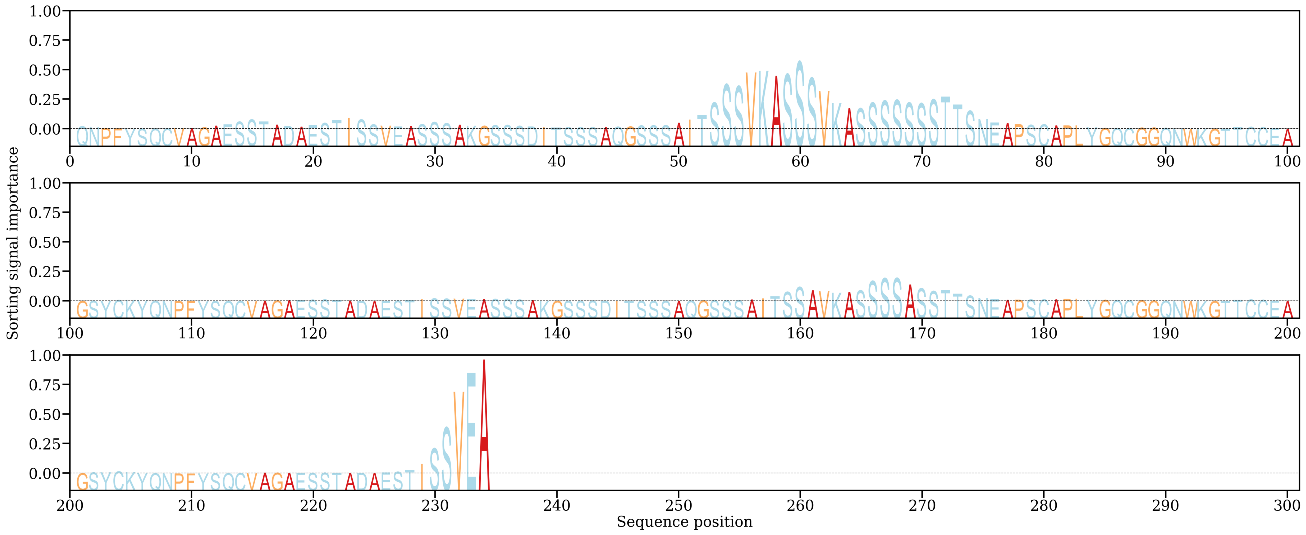


**Starmera_pachycereana_scaf50**
**Predicted localizations:** Extracellular
**Predicted signals:** Signal peptide

| Localization | Cytoplasm | Nucleus | Extracellular | Cell membrane | Mitochondrion | Plastid | Endoplasmic reticulum | Lysosome/Vacuole | Golgi apparatus | Peroxisome |
|---|---|---|---|---|---|---|---|---|---|---|
| **Probability** | 0.2283 | 0.1475 | 0.9320 | 0.1755 | 0.0597 | 0.0037 | 0.1944 | 0.0769 | 0.0856 | 0.0011 |

**Sorting Signal Importance. Donwload:** PNG (/services/DeepLoc-2.0/tmp/64A6C5DD00002D2A7A92F90C/alpha_starmera_pachycereana_scaf50.png) / CSV (/services/DeepLoc-2.0/tmp/64A6C5DD00002D2A7A92F90C/alpha_starmera_pachycereana_scaf50.csv)

Starmera_pachycereana_scaf50
Predicted Signals: Signal peptide

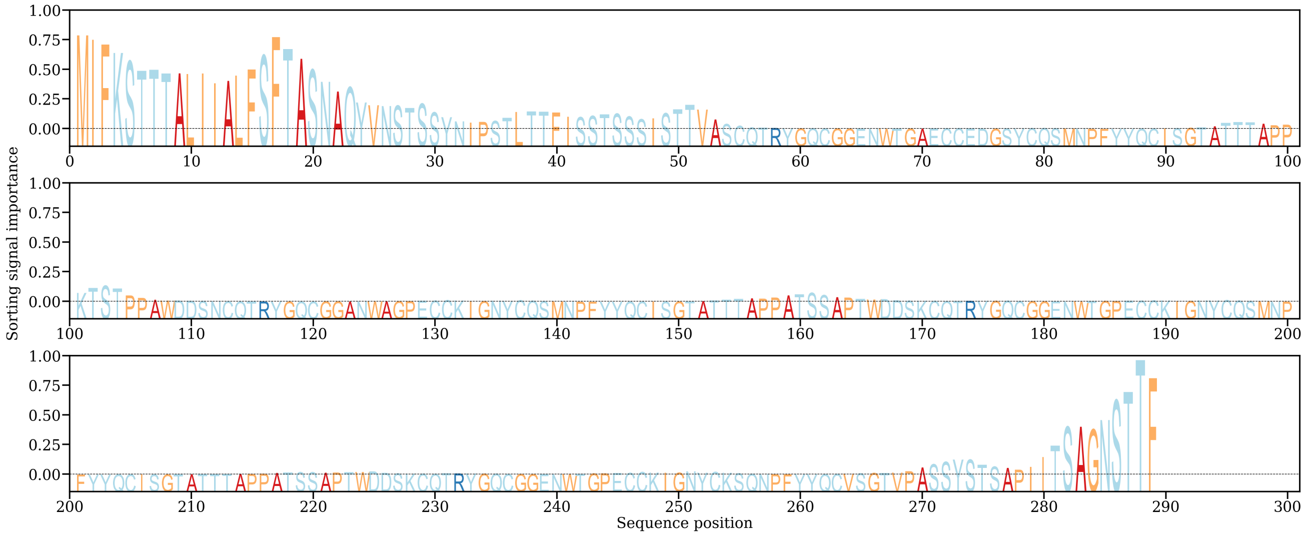


**Starmera_amethionina_scaf133**
**Predicted localizations:** Extracellular
**Predicted signals:** Signal peptide

| Localization | Cytoplasm | Nucleus | Extracellular | Cell membrane | Mitochondrion | Plastid | Endoplasmic reticulum | Lysosome/Vacuole | Golgi apparatus | Peroxisome |
|---|---|---|---|---|---|---|---|---|---|---|
| **Probability** | 0.1705 | 0.0761 | 0.9473 | 0.1268 | 0.0496 | 0.0024 | 0.3363 | 0.2836 | 0.4312 | 0.0026 |

**Sorting Signal Importance. Donwload:** PNG (/services/DeepLoc-2.0/tmp/64A6C5DD00002D2A7A92F90C/alpha_starmera_amethionina_scaf133.png) / CSV (/services/DeepLoc-2.0/tmp/64A6C5DD00002D2A7A92F90C/alpha_starmera_amethionina_scaf133.csv)

Starmera_amethionina_scaf133
Predicted Signals: Signal peptide

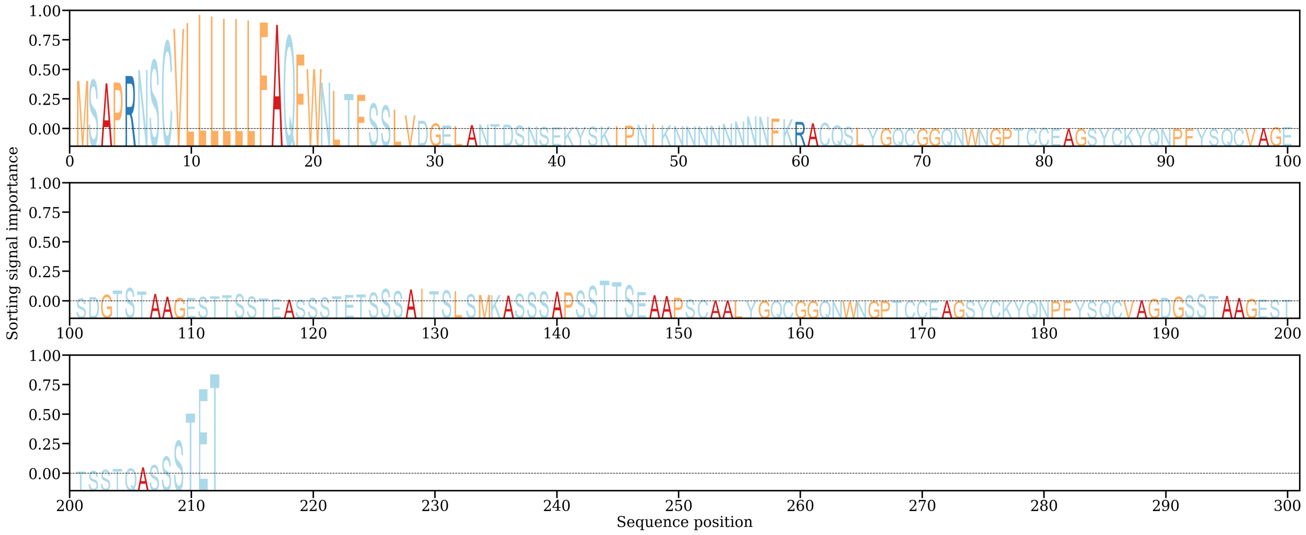


**Starmera_amethionina_scaf118**
**Predicted localizations:** Extracellular
**Predicted signals:**

| Localization | Cytoplasm | Nucleus | Extracellular | Cell membrane | Mitochondrion | Plastid | Endoplasmic reticulum | Lysosome/Vacuole | Golgi apparatus | Peroxisome |
|---|---|---|---|---|---|---|---|---|---|---|
| **Probability** | 0.2933 | 0.3248 | 0.8381 | 0.0992 | 0.0324 | 0.0017 | 0.1071 | 0.0410 | 0.0603 | 0.0292 |

**Sorting Signal Importance. Donwload:** PNG (/services/DeepLoc-2.0/tmp/64A6C5DD00002D2A7A92F90C/alpha_starmera_amethionina_scaf118.png) **/** CSV (/services/DeepLoc-2.0/tmp/64A6C5DD00002D2A7A92F90C/alpha_starmera_amethionina_scaf118.csv)

Starmera_amethionina_scaf118
Predicted Signals:

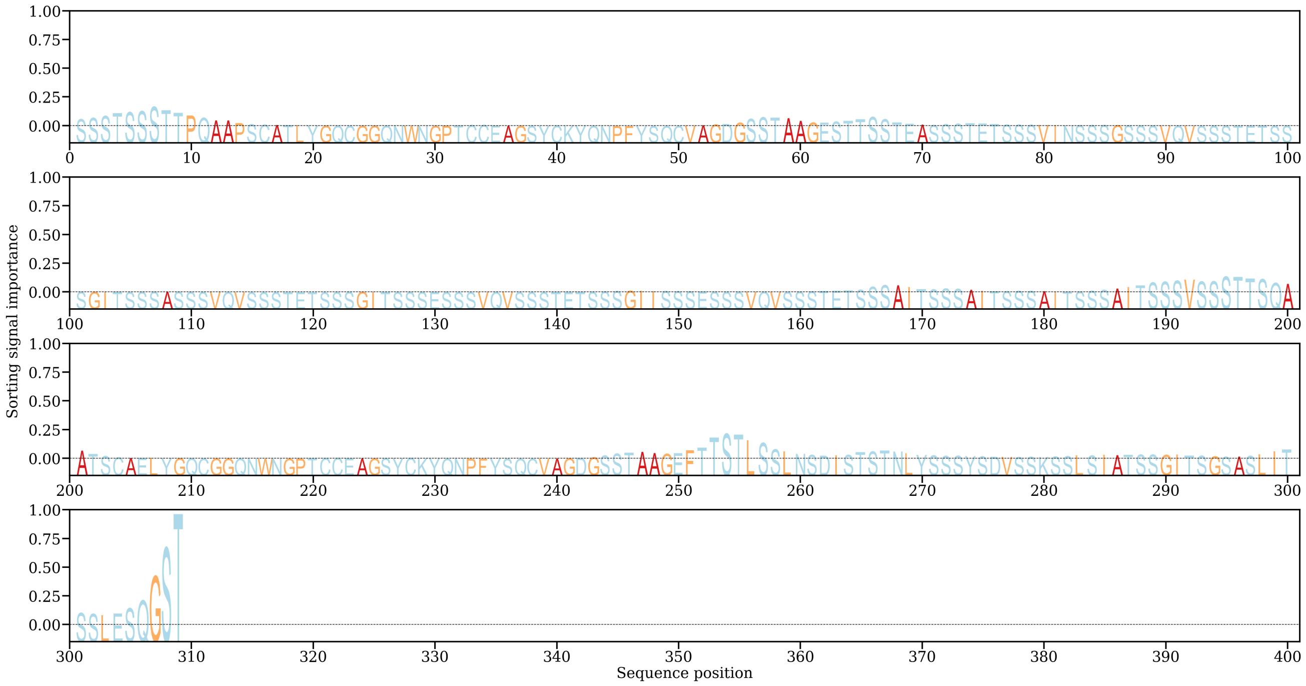


**Starmera_amethionina_scaf68**
**Predicted localizations:** Extracellular
**Predicted signals:**

| Localization | Cytoplasm | Nucleus | Extracellular | Cell membrane | Mitochondrion | Plastid | Endoplasmic reticulum | Lysosome/Vacuole | Golgi apparatus | Peroxisome |
|---|---|---|---|---|---|---|---|---|---|---|
| **Probability** | 0.4737 | 0.1575 | 0.6429 | 0.1127 | 0.0452 | 0.0034 | 0.0641 | 0.0679 | 0.0525 | 0.0133 |

**Sorting Signal Importance. Donwload:** PNG (/services/DeepLoc-2.0/tmp/64A6C5DD00002D2A7A92F90C/alpha_starmera_amethionina_scaf68.png) **/** CSV (/services/DeepLoc-2.0/tmp/64A6C5DD00002D2A7A92F90C/alpha_starmera_amethionina_scaf68.csv)

Starmera_amethionina_scaf68
Predicted Signals:

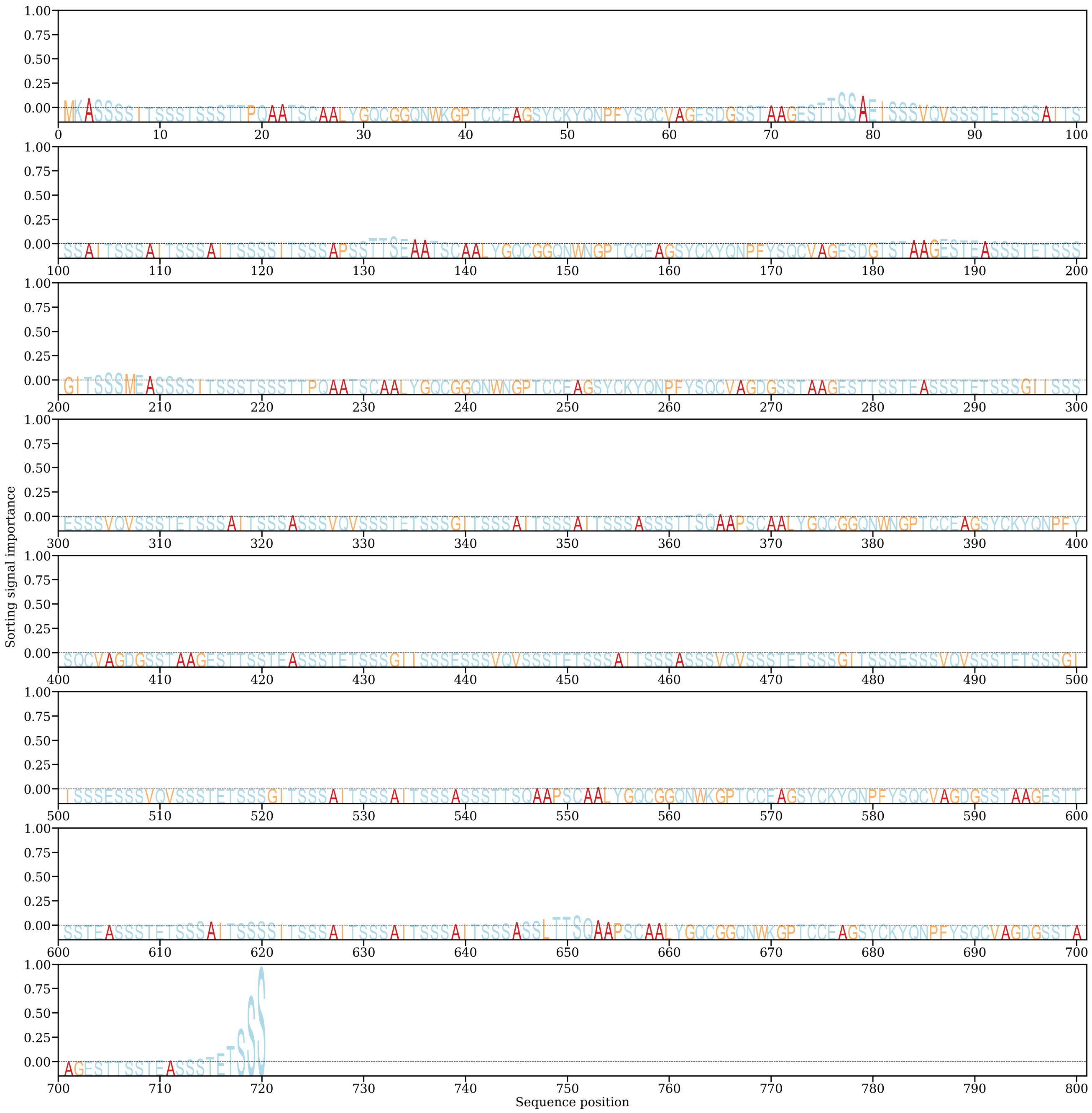


**Starmera_amethionina_scaf61**
**Predicted localizations:** Extracellular
**Predicted signals:** Signal peptide

| Localization | Cytoplasm | Nucleus | Extracellular | Cell membrane | Mitochondrion | Plastid | Endoplasmic reticulum | Lysosome/Vacuole | Golgi apparatus | Peroxisome |
|---|---|---|---|---|---|---|---|---|---|---|
| **Probability** | 0.2169 | 0.1439 | 0.9496 | 0.1919 | 0.0519 | 0.0052 | 0.1253 | 0.0561 | 0.0753 | 0.0010 |

**Sorting Signal Importance. Donwload:** PNG (/services/DeepLoc-2.0/tmp/64A6C5DD00002D2A7A92F90C/alpha_starmera_amethionina_scaf61.png) **/** CSV (/services/DeepLoc-2.0/tmp/64A6C5DD00002D2A7A92F90C/alpha_starmera_amethionina_scaf61.csv)

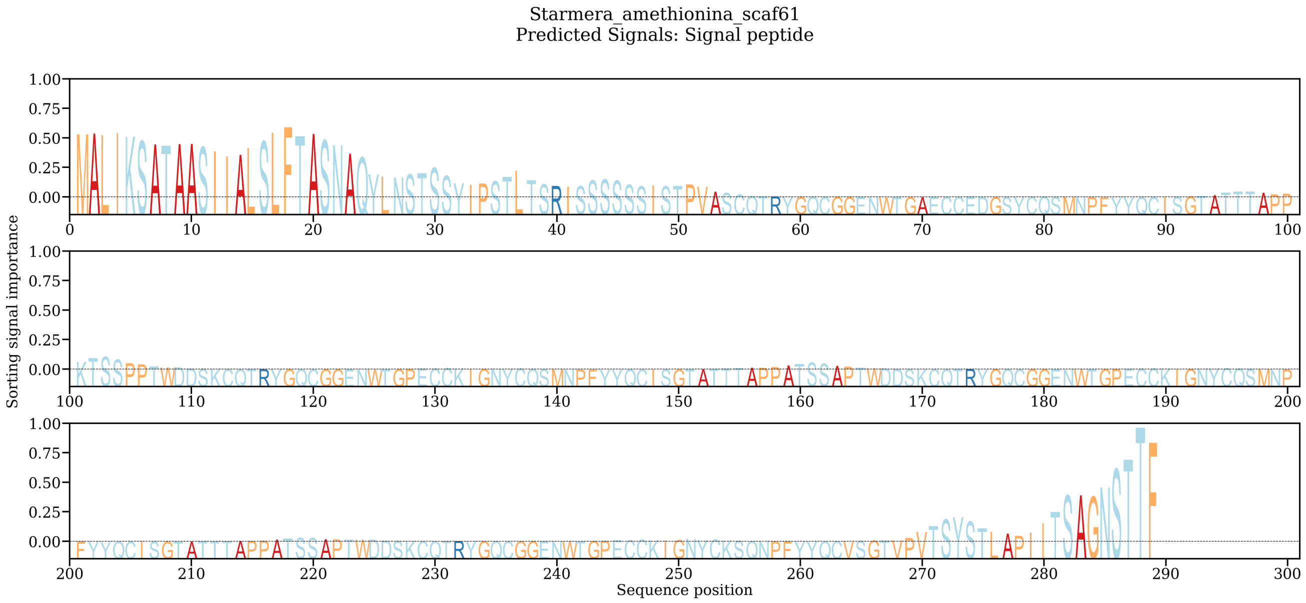


**Starmera_caribaea_scaf1610**

**Predicted localizations:** Extracellular

**Predicted signals:** Signal peptide

| Localization | Cytoplasm | Nucleus | Extracellular | Cell membrane | Mitochondrion | Plastid | Endoplasmic reticulum | Lysosome/Vacuole | Golgi apparatus | Peroxisome |
|---|---|---|---|---|---|---|---|---|---|---|
| **Probability** | 0.2977 | 0.2779 | 0.9735 | 0.1091 | 0.0334 | 0.0024 | 0.0552 | 0.0447 | 0.0642 | 0.0138 |

**Sorting Signal Importance. Donwload:** PNG (/services/DeepLoc-2.0/tmp/64A6C5DD00002D2A7A92F90C/alpha_starmera_caribaea_scaf1610.png) **/** CSV (/services/DeepLoc-2.0/tmp/64A6C5DD00002D2A7A92F90C/alpha_starmera_caribaea_scaf1610.csv)

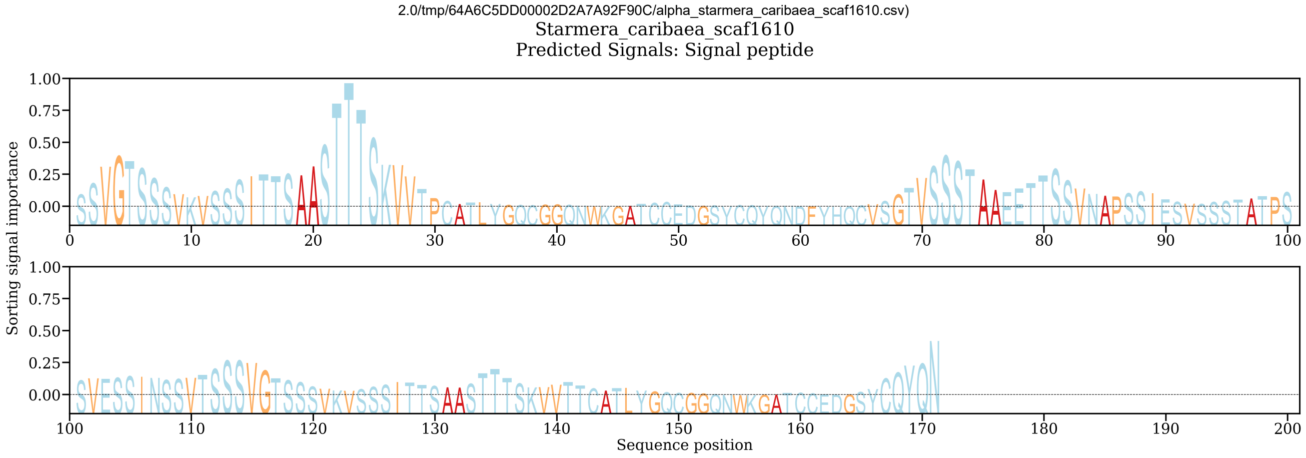


**Starmera_caribaea_scaf698**

**Predicted localizations:** Extracellular

**Predicted signals:** Signal peptide

| Localization | Cytoplasm | Nucleus | Extracellular | Cell membrane | Mitochondrion | Plastid | Endoplasmic reticulum | Lysosome/Vacuole | Golgi apparatus | Peroxisome |
|---|---|---|---|---|---|---|---|---|---|---|
| **Probability** | 0.1918 | 0.0751 | 0.9662 | 0.1178 | 0.0877 | 0.0043 | 0.2374 | 0.1785 | 0.3116 | 0.0038 |

**Sorting Signal Importance. Donwload:** PNG (/services/DeepLoc-2.0/tmp/64A6C5DD00002D2A7A92F90C/alpha_starmera_caribaea_scaf698.png) **/** CSV (/services/DeepLoc-2.0/tmp/64A6C5DD00002D2A7A92F90C/alpha_starmera_caribaea_scaf698.csv)

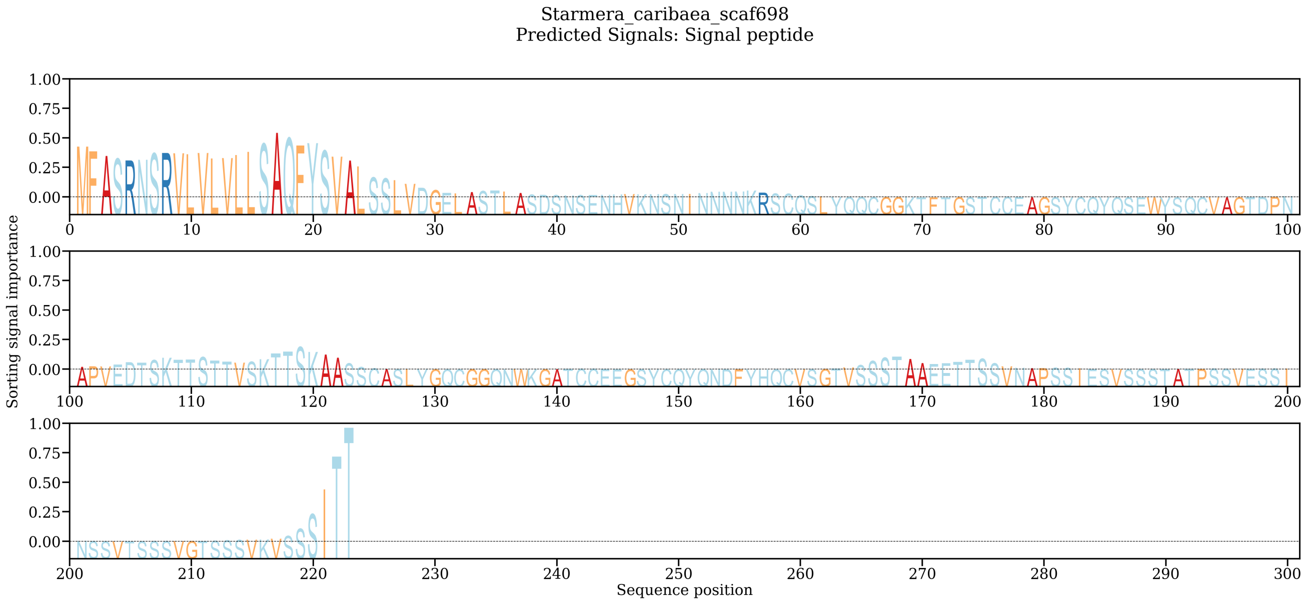


**Starmera_caribaea_scaf275**
**Predicted localizations:** Extracellular
**Predicted signals:**

| Localization | Cytoplasm | Nucleus | Extracellular | Cell membrane | Mitochondrion | Plastid | Endoplasmic reticulum | Lysosome/Vacuole | Golgi apparatus | Peroxisome |
|---|---|---|---|---|---|---|---|---|---|---|
| **Probability** | 0.2385 | 0.2818 | 0.9052 | 0.1967 | 0.0203 | 0.0028 | 0.1332 | 0.0312 | 0.1012 | 0.0351 |

**Sorting Signal Importance. Donwload:** PNG (/services/DeepLoc-2.0/tmp/64A6C5DD00002D2A7A92F90C/alpha_starmera_caribaea_scaf275.png) / CSV (/services/DeepLoc-2.0/tmp/64A6C5DD00002D2A7A92F90C/alpha_starmera_caribaea_scaf275.csv)

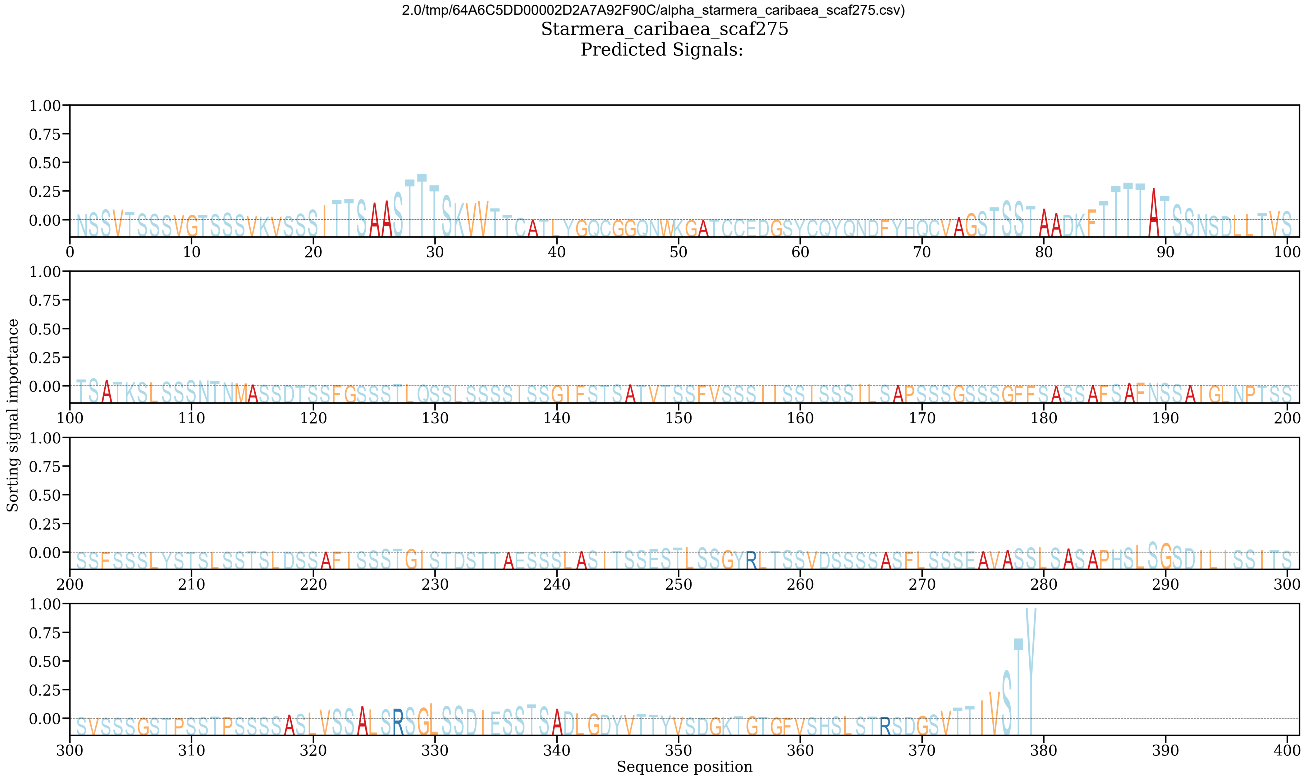


**Starmera_caribaea_scaf1564**
**Predicted localizations:** Extracellular
**Predicted signals:** Signal peptide

| Localization | Cytoplasm | Nucleus | Extracellular | Cell membrane | Mitochondrion | Plastid | Endoplasmic reticulum | Lysosome/Vacuole | Golgi apparatus | Peroxisome |
|---|---|---|---|---|---|---|---|---|---|---|
| **Probability** | 0.3357 | 0.2625 | 0.8324 | 0.0596 | 0.0222 | 0.0023 | 0.1041 | 0.0883 | 0.0880 | 0.0489 |

**Sorting Signal Importance. Donwload:** PNG (/services/DeepLoc-2.0/tmp/64A6C5DD00002D2A7A92F90C/alpha_starmera_caribaea_scaf1564.png) / CSV (/services/DeepLoc-2.0/tmp/64A6C5DD00002D2A7A92F90C/alpha_starmera_caribaea_scaf1564.csv)

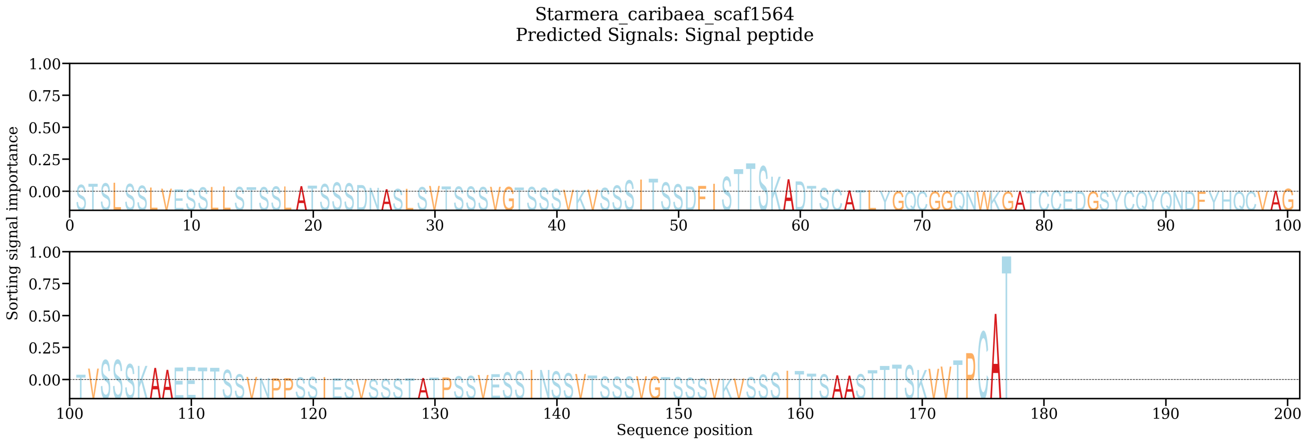


##### Starmera_caribaea_scaf1582

**Predicted localizations:** Extracellular

**Predicted signals:**

| Localization | Cytoplasm | Nucleus | Extracellular | Cell membrane | Mitochondrion | Plastid | Endoplasmic reticulum | Lysosome/Vacuole | Golgi apparatus | Peroxisome |
|---|---|---|---|---|---|---|---|---|---|---|
| **Probability** | 0.2956 | 0.2649 | 0.9047 | 0.0666 | 0.0563 | 0.0016 | 0.0793 | 0.0649 | 0.0585 | 0.0765 |

**Sorting Signal Importance. Donwload:** PNG (/services/DeepLoc-2.0/tmp/64A6C5DD00002D2A7A92F90C/alpha_starmera_caribaea_scaf1582.png) **/** CSV (/services/DeepLoc-2.0/tmp/64A6C5DD00002D2A7A92F90C/alpha_starmera_caribaea_scaf1582.csv)

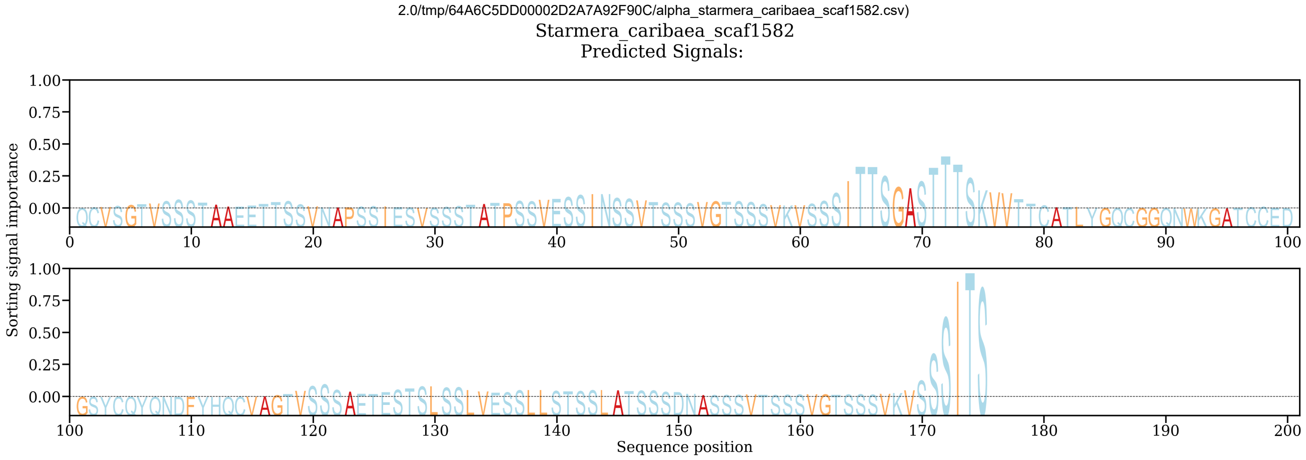


##### Starmera_caribaea_scaf29

**Predicted localizations:** Extracellular

**Predicted signals:** Signal peptide

| Localization | Cytoplasm | Nucleus | Extracellular | Cell membrane | Mitochondrion | Plastid | Endoplasmic reticulum | Lysosome/Vacuole | Golgi apparatus | Peroxisome |
|---|---|---|---|---|---|---|---|---|---|---|
| **Probability** | 0.2121 | 0.1452 | 0.9356 | 0.1653 | 0.0543 | 0.0047 | 0.1512 | 0.0718 | 0.0715 | 0.0007 |

**Sorting Signal Importance. Donwload:** PNG (/services/DeepLoc-2.0/tmp/64A6C5DD00002D2A7A92F90C/alpha_starmera_caribaea_scaf29.png) **/** CSV (/services/DeepLoc-2.0/tmp/64A6C5DD00002D2A7A92F90C/alpha_starmera_caribaea_scaf29.csv)

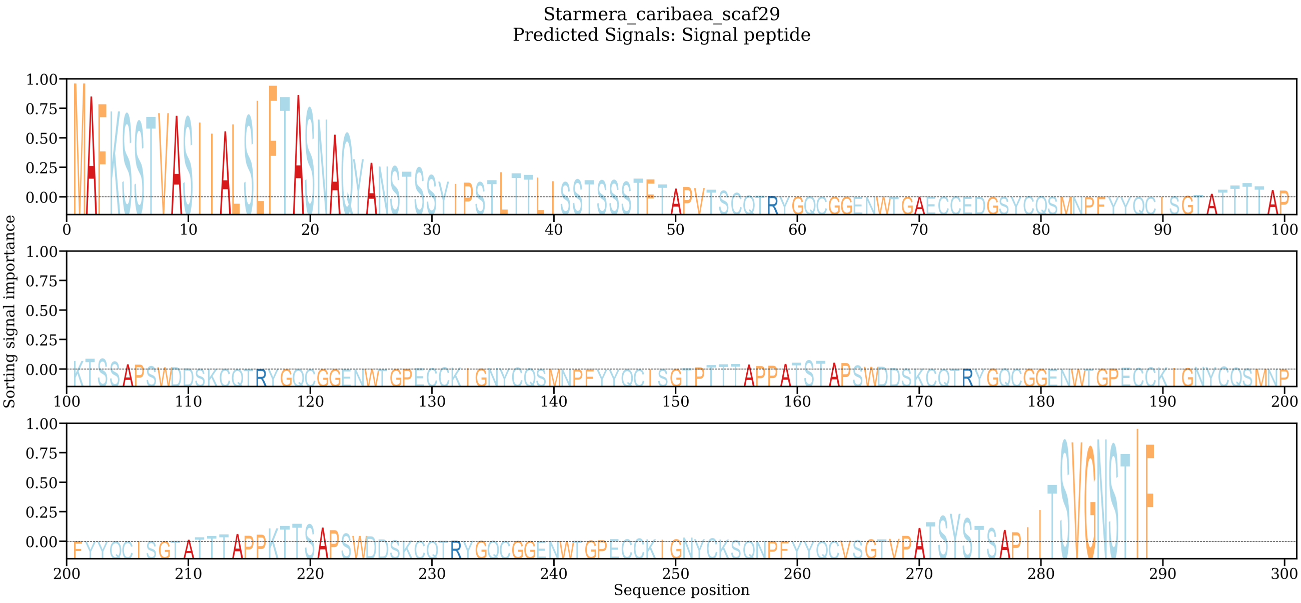


**Starmera_stellimalicola_scaf11**
**Predicted localizations:** Extracellular
**Predicted signals:** Signal peptide

| Localization | Cytoplasm | Nucleus | Extracellular | Cell membrane | Mitochondrion | Plastid | Endoplasmic reticulum | Lysosome/Vacuole | Golgi apparatus | Peroxisome |
|---|---|---|---|---|---|---|---|---|---|---|
| **Probability** | 0.1914 | 0.1837 | 0.7528 | 0.1077 | 0.0773 | 0.0026 | 0.0527 | 0.1749 | 0.1123 | 0.0221 |

**Sorting Signal Importance. Donwload:** PNG (/services/DeepLoc-2.0/tmp/64A6C5DD00002D2A7A92F90C/alpha_starmera_stellimalicola_scaf11.png) **/** CSV (/services/DeepLoc-2.0/tmp/64A6C5DD00002D2A7A92F90C/alpha_starmera_stellimalicola_scaf11.csv)

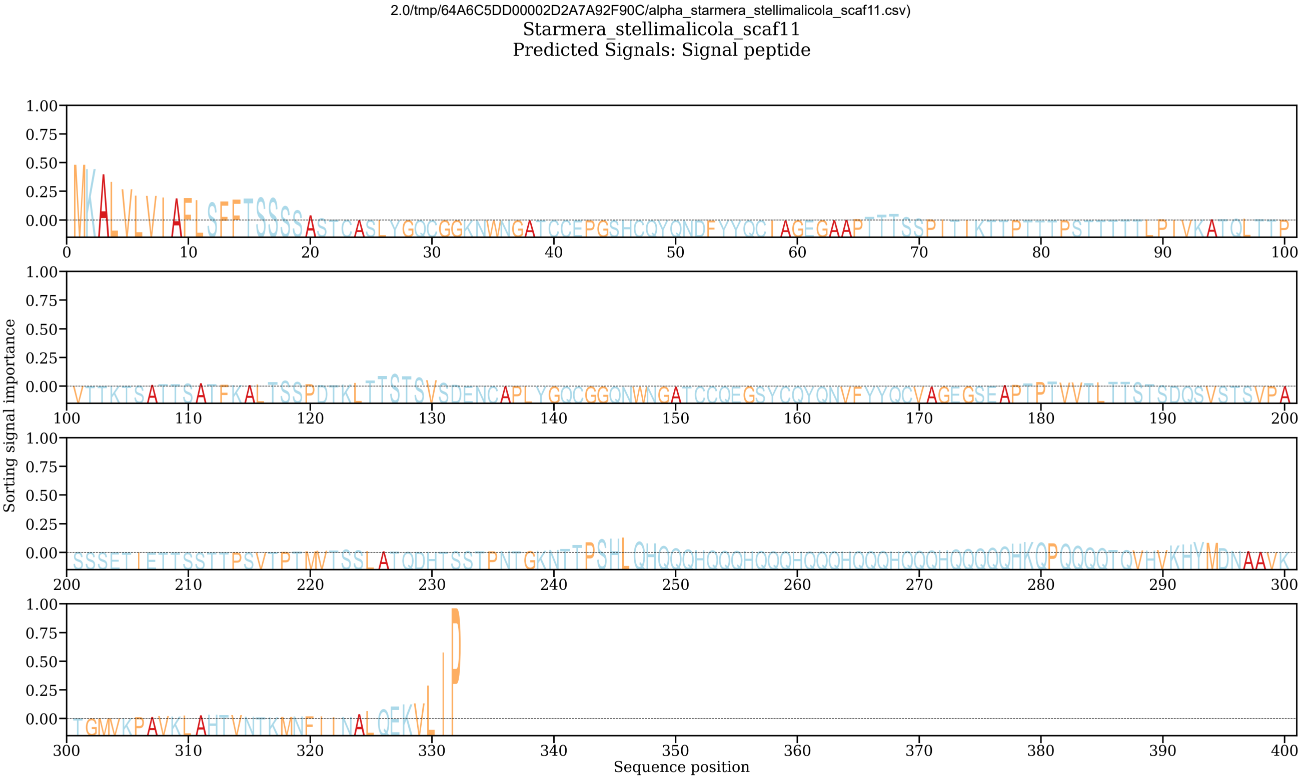


**Starmera_stellimalicola_scaf15**
**Predicted localizations:** Extracellular
**Predicted signals:** Signal peptide

| Localization | Cytoplasm | Nucleus | Extracellular | Cell membrane | Mitochondrion | Plastid | Endoplasmic reticulum | Lysosome/Vacuole | Golgi apparatus | Peroxisome |
|---|---|---|---|---|---|---|---|---|---|---|
| **Probability** | 0.1058 | 0.0728 | 0.9517 | 0.0854 | 0.0302 | 0.0134 | 0.0633 | 0.0932 | 0.0693 | 0.0012 |

**Sorting Signal Importance. Donwload:** PNG (/services/DeepLoc-2.0/tmp/64A6C5DD00002D2A7A92F90C/alpha_starmera_stellimalicola_scaf15.png) **/** CSV (/services/DeepLoc-2.0/tmp/64A6C5DD00002D2A7A92F90C/alpha_starmera_stellimalicola_scaf15.csv)

Starmera_stellimalicola_scaf15
Predicted Signals: Signal peptide

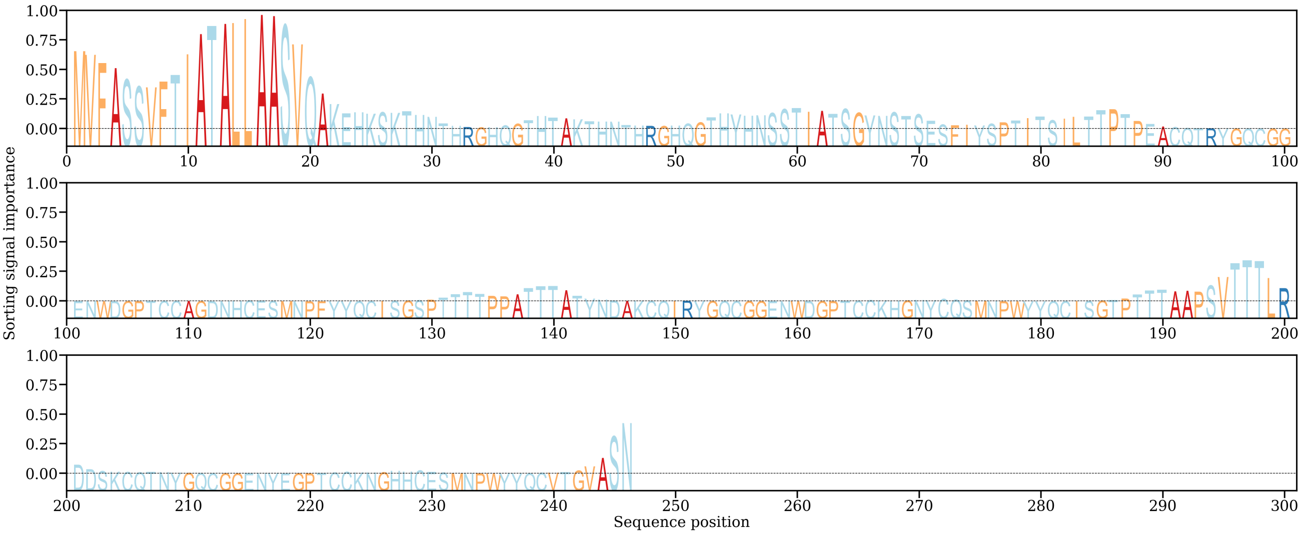


Found device of type cuda: Tesla V100-PCIE-16GB Loading model... Running model on device of type cuda: Tesla V100-PCIE-16GB Model loaded in 45.69s Attention plotted in 74.26s Model run time: 1.34s Finished prediction in 124.66s

**Fig. S3**

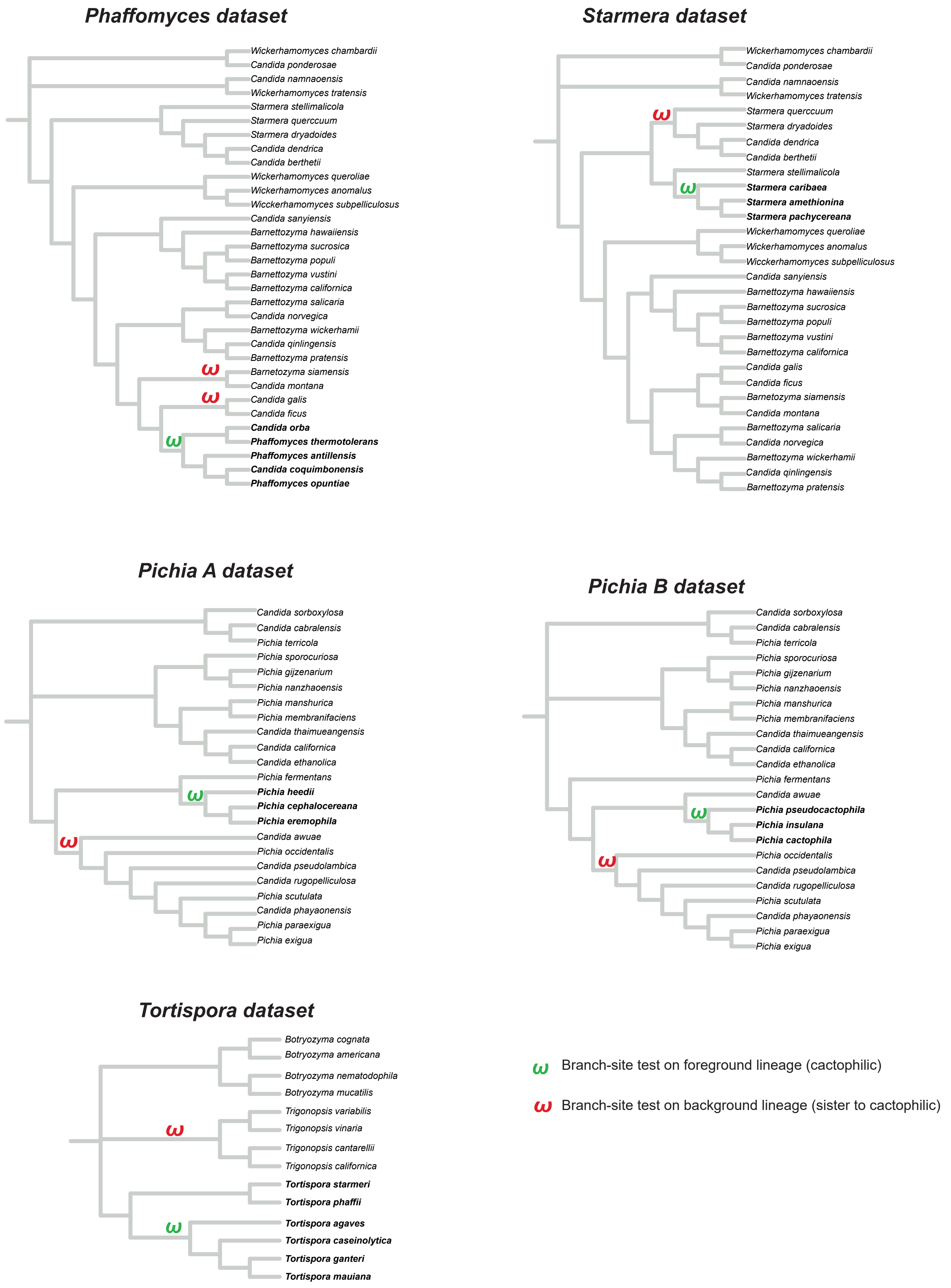
Phaffomyces dataset
Wickerhamomyces chambardii
Candida ponderosae
Candida namnaoensis
Wickerhamomyces tratensis
Starmera stellimalicola
Starmera quercuum
Starmera dryadoides
Candida dendrica
Candida berthetii
Wickerhamomyces queroliae
Wickerhamomyces anomalus
Wicckerhamomyces subpelliculosus
Candida sanyiensis
Barnettozyma hawaiiensis
Barnettozyma sucrosica
Barnettozyma populi
Barnettozyma vustini
Barnettozyma californica
Barnettozyma salicaria
Candida norvegica
Barnettozyma wickerhamii
Candida qinlingensis
Barnettozyma pratensis
Barnetozyma siamensis
Candida montana
Candida galis
Candida ficus
Candida orba
Phaffomyces thermotolerans
Phaffomyces antillensis
Candida coquimbonensis
Phaffomyces opuntiae
Starmera dataset
Wickerhamomyces chambardii
Candida ponderosae
Candida namnaoensis
Wickerhamomyces tratensis
Starmera quercuum
Starmera dryadoides
Candida dendrica
Candida berthetii
Starmera stellimalicola
Starmera caribaea
Starmera amethionina
Starmera pachycereana
Wickerhamomyces queroliae
Wickerhamomyces anomalus
Wicckerhamomyces subpelliculosus
Candida sanyiensis
Barnettozyma hawaiiensis
Barnettozyma sucrosica
Barnettozyma populi
Barnettozyma vustini
Barnettozyma californica
Candida galis
Candida ficus
Barnetozyma siamensis
Candida montana
Barnettozyma salicaria
Candida norvegica
Barnettozyma wickerhamii
Candida qinlingensis
Barnettozyma pratensis
Pichia A dataset
Candida sorboxylosa
Candida cabralensis
Pichia terricola
Pichia sporocuriosa
Pichia gijzenarium
Pichia nanzhaoensis
Pichia manshurica
Pichia membranifaciens
Candida thaimueangensis
Candida californica
Candida ethanolica
Pichia fermentans
Pichia heedii
Pichia cephalocereana
Pichia eremophila
Candida awuae
Pichia occidentalis
Candida pseudolambica
Candida rugopelliculosa
Pichia scutulata
Candida phayaonensis
Pichia paraexigua
Pichia exigua
Pichia B dataset
Candida sorboxylosa
Candida cabralensis
Pichia terricola
Pichia sporocuriosa
Pichia gijzenarium
Pichia nanzhaoensis
Pichia manshurica
Pichia membranifaciens
Candida thaimueangensis
Candida californica
Candida ethanolica
Pichia fermentans
Candida awuae
Pichia pseudocactophila
Pichia insulana
Pichia cactophila
Pichia occidentalis
Candida pseudolambica
Candida rugopelliculosa
Pichia scutulata
Candida phayaonensis
Pichia paraexigua
Pichia exigua
Tortispora dataset
Botryozyma cognata
Botryozyma americana
Botryozyma nematodophila
Botryozyma mucatilis
Trigonopsis variabilis
Trigonopsis vinaria
Trigonopsis cantarellii
Trigonopsis californica
Tortispora starmeri
Tortispora phaffii
Tortispora agaves
Tortispora caseinolytica
Tortispora ganteri
Tortispora mauiana
ω Branch-site test on foreground lineage (cactophilic)
ω Branch-site test on background lineage (sister to cactophilic)

**Fig. S4**

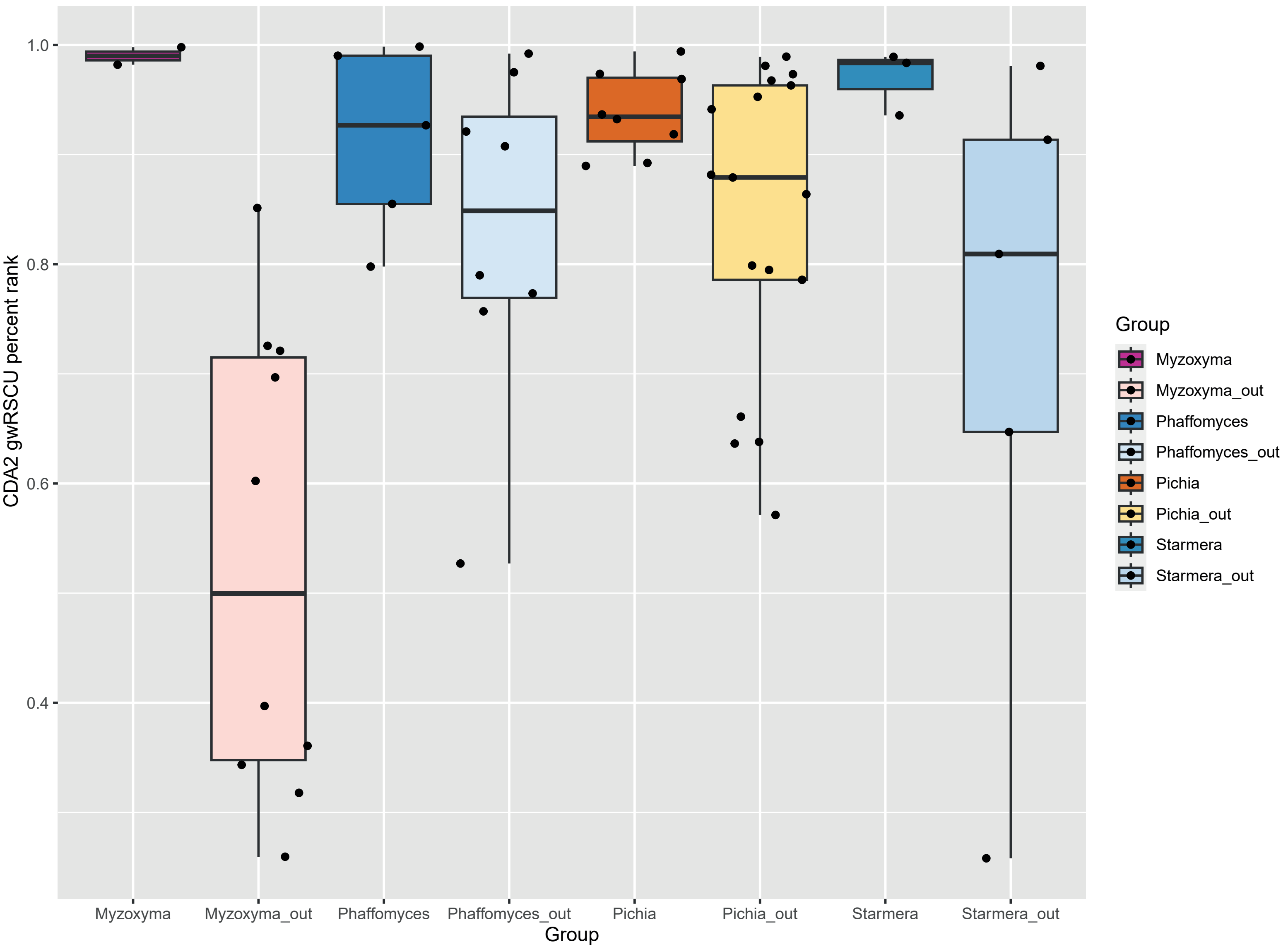

CDA2 gwRSCU percent rank
1.0
0.8
0.6
0.4
Myzoxyma
Myzoxyma_out
Phaffomyces
Phaffomyces_out
Pichia
Pichia_out
Starmera
Starmera_out
Group
Group
Myzoxyma
Myzoxyma_out
Phaffomyces
Phaffomyces_out
Pichia
Pichia_out
Starmera
Starmera_out

### Fig. S5

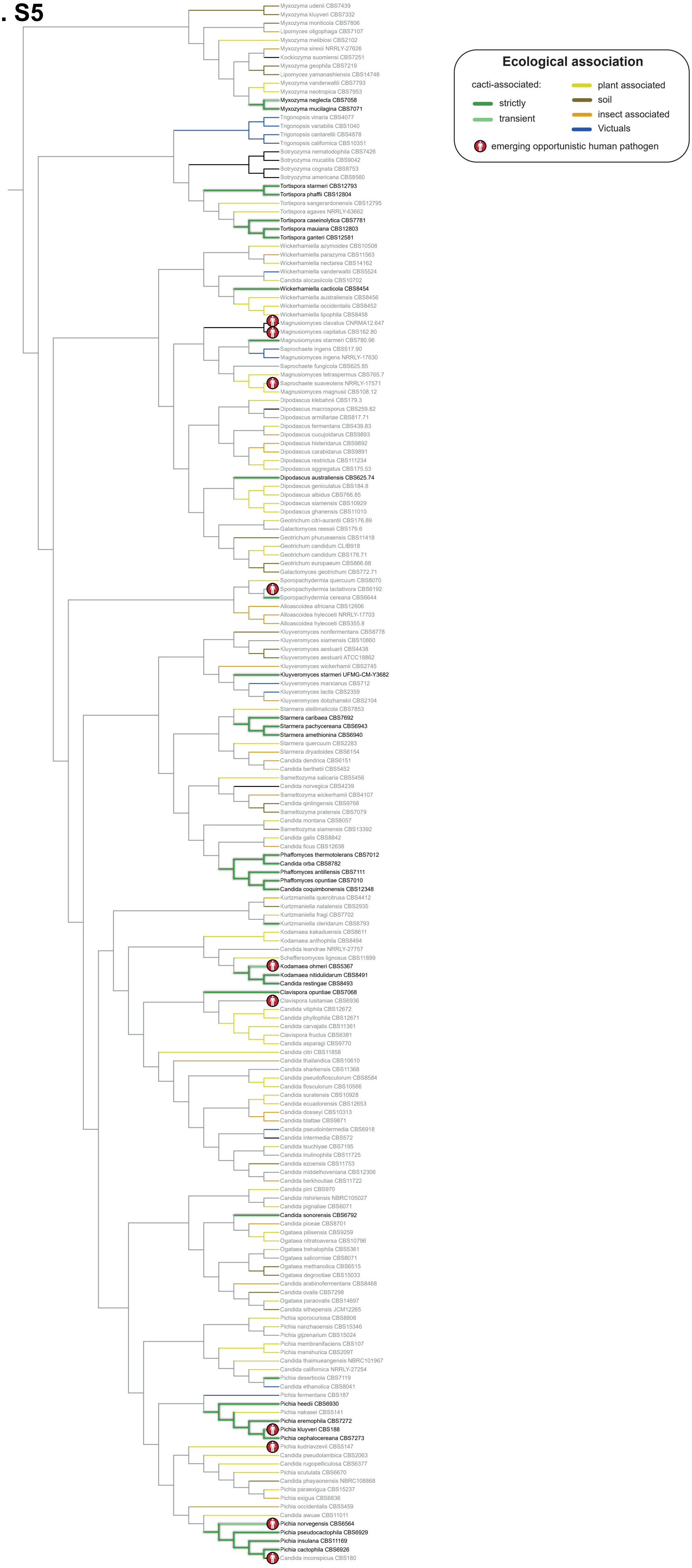

Ecological association
cacti-associated:
strictly
transient
plant associated
soil
insect associated
Victuals
emerging opportunistic human pathogen
Myxozyma udenii CBS7439
Myxozyma kluyveri CBS7332
Myxozyma monticola CBS7806
Lipomyces oligophaga CBS7107
Myxozyma melibiosi CBS2102
Myxozyma sirexii NRRLY-27626
Kockiozyma suomiensi CBS7251
Myxozyma geophila CBS7219
Lipomyces yamanashiensis CBS14748
Myxozyma vanderwaltii CBS7793
Myxozyma neotropica CBS7953
Myxozyma neglecta CBS7058
Myxozyma mucilagina CBS7071
Trigonopsis vinaria CBS4077
Trigonopsis variabilis CBS1040
Trigonopsis cantarellii CBS4878
Trigonopsis californica CBS10351
Botryozyma nematodophila CBS7426
Botryozyma mucatilis CBS9042
Botryozyma cognata CBS8753
Botryozyma americana CBS8560
Tortispora starmeri CBS12793
Tortispora phaffii CBS12804
Tortispora sangerardonensis CBS12795
Tortispora agaves NRRLY-63662
Tortispora caseinolytica CBS7781
Tortispora mauiana CBS12803
Tortispora ganteri CBS12581
Wickerhamiella azymoides CBS10508
Wickerhamiella parazyma CBS11563
Wickerhamiella nectarea CBS14162
Wickerhamiella vanderwaltii CBS5524
Candida alocasiicola CBS10702
Wickerhamiella cacticola CBS8454
Wickerhamiella australiensis CBS8456
Wickerhamiella occidentalis CBS8452
Wickerhamiella lipophila CBS8458
Magnusiomyces clavatus CNRMA12.647
Magnusiomyces capitatus CBS162.80
Magnusiomyces starmeri CBS780.96
Saprochaete ingens CBS517.90
Magnusiomyces ingens NRRLY-17630
Saprochaete fungicola CBS625.85
Magnusiomyces tetraspermus CBS765.7
Saprochaete suaveolens NRRLY-17571
Magnusiomyces magnusii CBS108.12
Dipodascus klebahnii CBS179.3
Dipodascus macrosporus CBS259.82
Dipodascus armillariae CBS817.71
Dipodascus fermentans CBS439.83
Dipodascus cucujoidarus CBS9893
Dipodascus histeridarus CBS9892
Dipodascus carabidarus CBS9891
Dipodascus restrictus CBS111234
Dipodascus aggregatus CBS175.53
Dipodascus australiensis CBS625.74
Dipodascus geniculatus CBS184.8
Dipodascus albidus CBS766.85
Dipodascus siamensis CBS10929
Dipodascus ghanensis CBS11010
Geotrichum citri-aurantii CBS176.89
Galactomyces reessii CBS179.6
Geotrichum phurueaensis CBS11418
Geotrichum candidum CLIB918
Geotrichum candidum CBS178.71
Geotrichum europaeum CBS866.68
Galactomyces geotrichum CBS772.71
Sporopachydermia quercuum CBS8070
Sporopachydermia lactativora CBS6192
Sporopachydermia cereana CBS6644
Alloascoidea africana CBS12606
Alloascoidea hylecoeti NRRLY-17703
Alloascoidea hylecoeti CBS355.8
Kluyveromyces nonfermentans CBS8778
Kluyveromyces siamensis CBS10860
Kluyveromyces aestuarii CBS4438
Kluyveromyces aestuarii ATCC18862
Kluyveromyces wickerhamii CBS2745
Kluyveromyces starmeri UFMG-CM-Y3682
Kluyveromyces marxianus CBS712
Kluyveromyces lactis CBS2359
Kluyveromyces dobzhanskii CBS2104
Starmera stellimalicola CBS7853
Starmera caribaea CBS7692
Starmera pachycereana CBS6943
Starmera amethionina CBS6940
Starmera quercuum CBS2283
Starmera dryadoides CBS6154
Candida dendrica CBS6151
Candida berthetii CBS5452
Barnettozyma salicaria CBS5456
Candida norvegica CBS4239
Barnettozyma wickerhamii CBS4107
Candida qinlingensis CBS9768
Barnettozyma pratensis CBS7079
Candida montana CBS8057
Barnettozyma siamensis CBS13392
Candida galis CBS8842
Candida ficus CBS12638
Phaffomyces thermotolerans CBS7012
Candida orba CBS8782
Phaffomyces antillensis CBS7111
Phaffomyces opuntiae CBS7010
Candida coquimbonensis CBS12348
Kurtzmaniella quercitrusa CBS4412
Kurtzmaniella natalensis CBS2935
Kurtzmaniella fragi CBS7702
Kurtzmaniella cleridarum CBS8793
Kodamaea kakaduensis CBS8611
Kodamaea anthophila CBS8494
Candida leandrae NRRLY-27757
Scheffersomyces lignosus CBS11899
Kodamaea ohmeri CBS5367
Kodamaea nitidulidarum CBS8491
Candida restingae CBS8493
Clavispora opuntiae CBS7068
Clavispora lusitaniae CBS6936
Candida vitiphila CBS12672
Candida phyllophila CBS12671
Candida carvajalis CBS11361
Clavispora fructus CBS6381
Candida asparagi CBS9770
Candida citri CBS11858
Candida thailandica CBS10610
Candida sharkensis CBS11368
Candida pseudoflosculorum CBS8584
Candida flosculorum CBS10566
Candida suratensis CBS10928
Candida ecuadorensis CBS12653
Candida dosseyi CBS10313
Candida blattae CBS9871
Candida pseudointermedia CBS6918
Candida intermedia CBS572
Candida tsuchiyae CBS7195
Candida inulinophila CBS11725
Candida ezoensis CBS11753
Candida middelhoveniana CBS12306
Candida berkhoutiae CBS11722
Candida pini CBS970
Candida rishiriensis NBRC105027
Candida pignaliae CBS6071
Candida sonorensis CBS6792
Candida piceae CBS8701
Ogataea pilisensis CBS9259
Ogataea nitratoaversa CBS10796
Ogataea trehalophila CBS5361
Ogataea salicorniae CBS8071
Ogataea methanolica CBS6515
Ogataea degrootiae CBS15033
Candida arabinofermentans CBS8468
Candida ovalis CBS7298
Ogataea paraovalis CBS14697
Candida sithepensis JCM12265
Pichia sporocuriosa CBS8806
Pichia nanzhaoensis CBS15346
Pichia gijzenarium CBS15024
Pichia membranifaciens CBS107
Pichia manshurica CBS209T
Candida thaimueangensis NBRC101967
Candida californica NRRLY-27254
Pichia deserticola CBS7119
Candida ethanolica CBS8041
Pichia fermentans CBS187
Pichia heedii CBS6930
Pichia nakasei CBS5141
Pichia eremophila CBS7272
Pichia kluyveri CBS188
Pichia cephalocereana CBS7273
Pichia kudriavzevii CBS5147
Candida pseudolambica CBS2063
Candida rugopelliculosa CBS6377
Pichia scutulata CBS6670
Candida phayaonensis NBRC108868
Pichia paraexigua CBS15237
Pichia exigua CBS6836
Pichia occidentalis CBS5459
Candida awuae CBS11011
Pichia norvegensis CBS6564
Pichia pseudocactophila CBS6929
Pichia insulana CBS11169
Pichia cactophila CBS6926
Candida inconspicua CBS180

Fig. S6

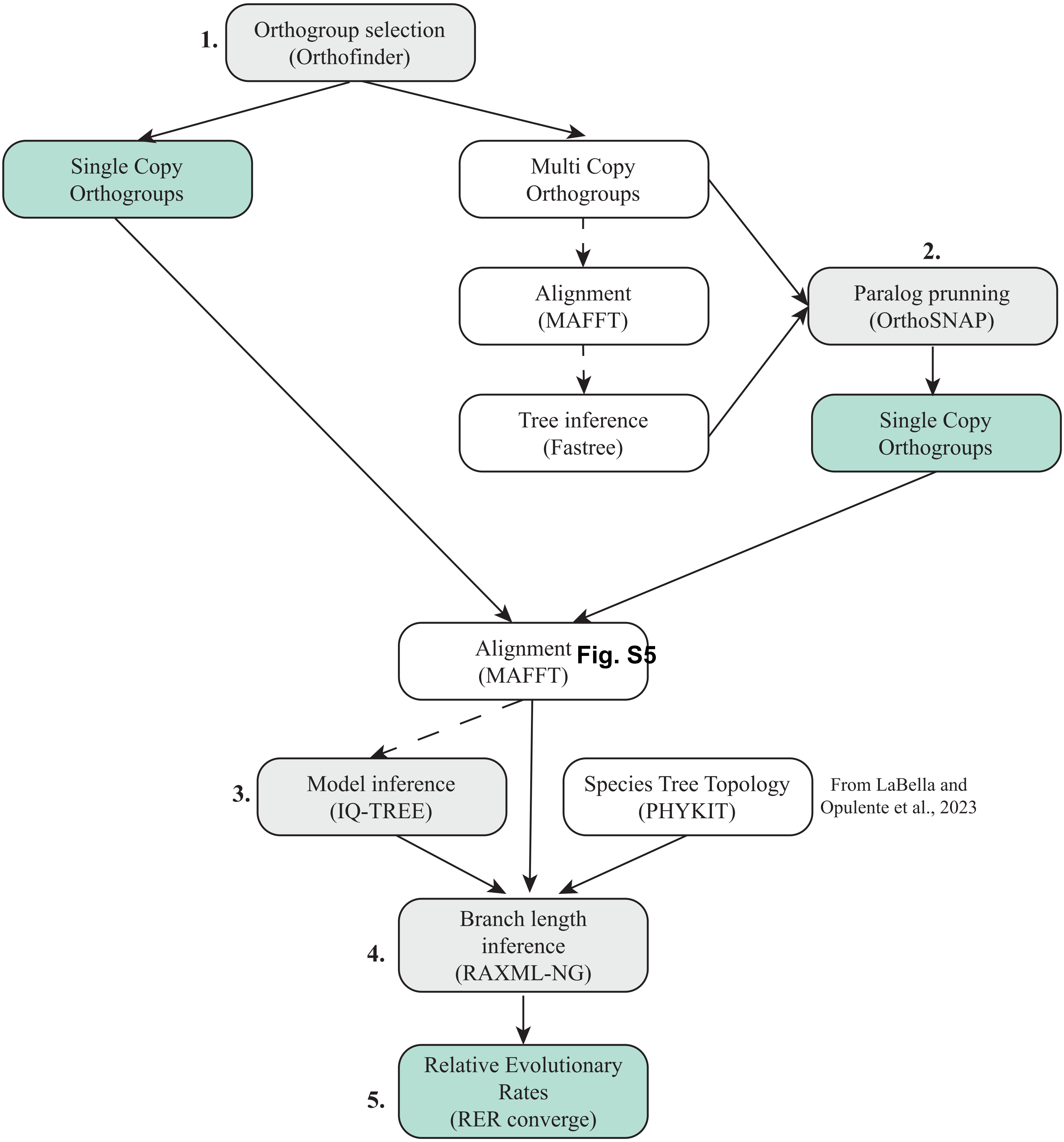

1.
Orthogroup selection (Orthofinder)
Single Copy Orthogroups
Multi Copy Orthogroups
Alignment (MAFFT)
Tree inference (Fastree)
2.
Paralog prunning (OrthoSNAP)
Single Copy Orthogroups
Alignment (MAFFT)
Fig. S5
3.
Model inference (IQ-TREE)
Species Tree Topology (PHYKIT)
From LaBella and Opulente et al., 2023
4.
Branch length inference (RAXML-NG)
5.
Relative Evolutionary Rates (RER converge)
